# α-hemolysin polymorphisms in methicillin-resistant *Staphylococcus aureus* clinical isolates regulate ADAM10-dependent neutrophil IL-1β secretion

**DOI:** 10.64898/2025.12.13.694139

**Authors:** Karl Liboro, Jolynn T. Chau, James D. Begando, Serena Abbondante, Angela Lackner, Yan Sun, Michaela E. Marshall, Nghi Ly, Victor D. Johnson, George R. Dubyak, Michael Gilmore, Reginald McNulty, Camille Andre, Eric Pearlman

## Abstract

*Staphylococcus aureus* α-hemolysin (Hla) is a major virulence factor that utilizes cell surface ADAM10 to oligomerize and form a functional heptameric pore. We show here that Hla from strain USA300 is required to induce IL-1β secretion by neutrophils and to cause severe corneal disease in mice. We also demonstrate that in contrast to USA300 and other clonal complex 8 (CC8) methicillin resistant *S. aureus* (MRSA) isolated from the skin, CC5 Hla from corneas of infected patients have single nucleotide polymorphisms (SNP) that result in two amino acid substitutions, D208E (Asp-Glu) and I275T (Ile-Thr). Structural modeling predicts CC5 Hla self-assembly and altered binding to ADAM10 that is distinct from CC8 Hla. The ADAM10 inhibitor GI254023X blocked neutrophil IL-1β secretion induced by Hla-expressing CC8, but not by CC5 conditioned media, indicating that these Hla polymorphisms play an important role in Hla receptor binding and neutrophil IL-1β secretion, and affect corneal disease severity.

## INTRODUCTION

The rising incidence of infections caused by multi-drug-resistant bacteria is a major public health threat worldwide^1–3^. Methicillin resistant *Staphylococcus aureus* (MRSA) strains are now prevalent in hospital and community environments, with USA300 emerging as the most common MRSA clone in the USA^4–6^. While MRSA are major causes of sepsis and of lung and soft tissue infections, they are also an important cause of potentially blinding corneal infections (keratitis) ^7–9^.

*S. aureus* secretes multiple exotoxins that contribute to virulence, playing roles in cell cytotoxicity and tissue damage^10,11^. Among these is α-hemolysin (Hla), a heptameric toxin that forms pores in the plasma membrane, resulting in membrane permeability and cell lysis^12,13^. Hla is an important virulence factor in human disease and has been the target of antibody- and drug-based treatments^10,14,15^. Although Hla can insert into the plasma membrane at high concentrations, its activity is significantly enhanced through binding to cell surface ADAM10 (A Disintegrin and Metalloprotease 10), although ADAM10 enzymatic activity is not required for Hla binding^16^.

Studies on the genetic diversity of community-acquired MRSA (CA-MRSA) and hospital-acquired MRSA (HA-MRSA) have reported differences in the expression of virulence factors and antibiotic resistance^17–22^. Molecular epidemiological analyses place *S. aureus* isolates into multiple clonal complexes (CC) where similar sequence types, determined by multi-locus sequence typing (MLST), are grouped together^23^. Further, a 2020 study by Bispo, Gilmore *et al.* reported that MRSA corneal and peri-ocular skin infections were enriched in specific clonal complexes. In that study, CC8 isolates were primarily isolated from infected skin and soft-tissues around the eye whereas CC5 isolates were primarily isolated from infections of the cornea and conjunctiva^24^.

In the current study, we assessed the role of Hla in a murine model of corneal infection with the MRSA strain USA300-LAC and with CC5 and CC8 clinical isolates from the Bispo and Gilmore study. Following intrastromal infection with USA300, we found that neutrophils comprise >90% of the cellular infiltrate in corneas during early infection, and that disease severity and bacterial killing were dependent on neutrophil production of IL-1β. We also found that corneal disease severity, neutrophil recruitment, and IL-1β secretion in infected corneas are dependent on Hla production, and that IL-1β secretion by neutrophils *in vitro* is dependent on Hla activation of the NLRP3 inflammasome.

To assess the broader implications of these findings, we examined clinical isolates from corneas or ocular abscesses of infected patients, focusing on the most common MRSA lineages in the USA, clonal complex 5 (CC5) and CC8 (which includes USA300). Interestingly, several isolates did not express Hla protein during growth *in vitro*, even though genome analysis shows that all isolates have an intact *hla* gene. Clinical isolates that did not secrete Hla did not induce neutrophil IL-1β secretion and most did not induce corneal disease, behaving similarly to an α-hemolysin mutant (ΔHla).

We also report that whereas the HLA genomic sequence of all CC8 isolates was identical, all CC5 clinical isolates from patient corneas had multiple short nucleotide polymorphisms SNPs, and that two of these SNPs predicted amino acid substitutions D208E (Asp-Glu) and I275T (Ile-Thr). ChimeraX and AlphaFold software predicted that these substitutions alter the conformation of Hla and its binding to ADAM10. This prediction was supported by functional studies showing that ADAM10 inhibition impairs Hla-mediated neutrophil IL-1β secretion induced by Hla from CC8 but not CC5 clinical isolates. Together, these findings provide insight into biology of Hla activity, and on the overall virulence of MRSA.

## RESULTS

### Bacterial clearance and corneal disease severity in MRSA keratitis are dependent on IL-1β and α-hemolysin

To determine if IL-1β regulates corneal disease severity caused by MRSA, we injected 1x10^3^ log phase the common MRSA isolate USA300 directly into the corneal stroma of IL-1α^-/-^, IL-1β^-/-^, and IL-1α/β^-/-^ mice. After 24 h, we found that C57BL/6 and IL-1α^-/-^ corneas exhibited pronounced corneal opacification, whereas IL-1β^-/-^ and IL-1α/β^-/-^ corneas show less corneal opacity overall but have distinct punctate opacities in the cornea (**Fig 1A,B**). Infected IL-1β^-/-^and IL-1α/β^-/-^ corneas also have significantly higher numbers of viable bacteria(colony forming units, CFU) compared with infected C57BL/6 and IL-1α^-/-^ corneas (**Fig 1C**). These results indicate that IL-1β and not IL-1α plays a major role in regulating bacterial growth and disease severity during MRSA corneal infections.

**Figure 1.**
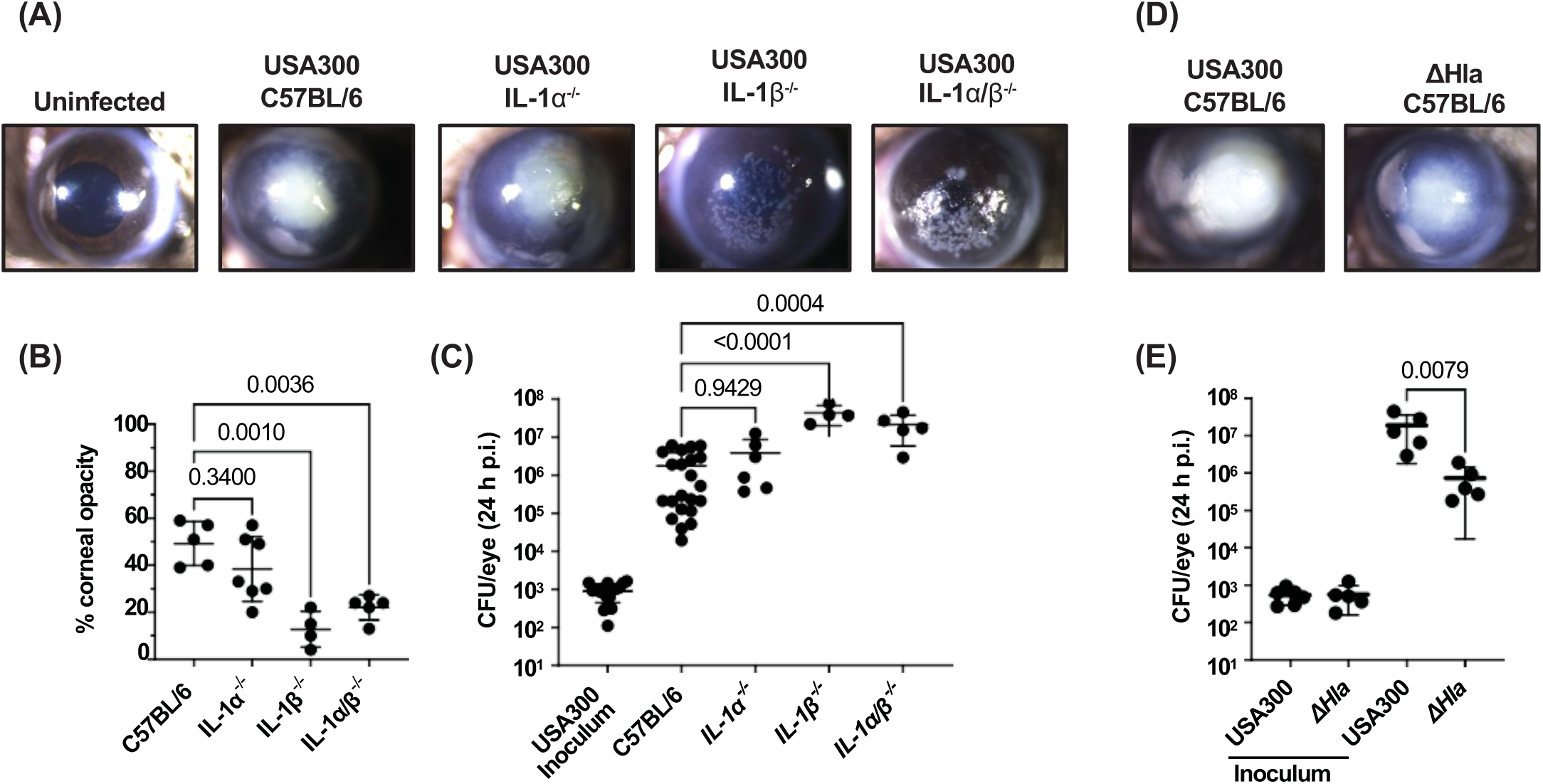
Bacterial killing and corneal disease severity in MRSA keratitis is dependent on IL-1β and α-hemolysin. 1x10^3^ CFU of USA300 or ΔHla were injected into corneal stroma of C57BL/6, IL-1α^-/-^, IL-1β^-/-^, or IL-1α/β^-/-^ mice and examined after 24 h. **A,D.** Representative images of infected corneas. **B.** Quantification of corneal opacification by image analysis. **C, E.** Quantification of viable bacteria by CFU. Statistical significance was determined by 1-way ANOVA followed by Dunnett’s multiple comparison test **(B, C)** or unpaired t-test **(E)**. Error bars represent mean +/- SD. Data points represent individual corneas.

To determine if Hla has a role in corneal disease severity, corneas were infected with USA300-LAC (termed USA300) or with an isogenic α-hemolysin transposon insertion mutant, NE1354^3–6^ (termed ΔHla) and examined as above. Results show two clear phenotypes where USA300 has pronounced, consolidated corneal opacification. In contrast, mice infected with ΔHla had less and more diffuse corneal opacification (**Fig 1D**) and significantly fewer viable bacteria (**Fig 1E**) than USA300 infected corneas. These data identify Hla as a critical virulence factor in bacterial survival in MRSA infected corneas, and are consistent with an earlier report using rabbit corneas infected with a laboratory strain of *S. aureus* ^25^

### Impaired neutrophil infiltration in infected IL-1β^-/-^ corneas result in bacterial replication in discrete pockets of the stroma

To characterize further the effect of IL-1β and Hla in MRSA infected corneas, whole eyes were paraffin embedded, sectioned, stained with H&E or with crystal violet to detect bacteria, or were incubated with antibodies to Ly6G to detect neutrophils.

Mammalian corneas are comprised of a multi-layered corneal epithelium, a single layer of corneal endothelial cells and the corneal stroma that comprises 80% of the cornea (**Fig 2A**). C57BL/6 corneas infected for 24 h with USA300 exhibited significant cellular infiltration, primarily comprised of Ly6G+ neutrophils (**Fig. 2B, upper panels)**. In contrast, ΔHla infected C57BL/6 and USA300 infected IL-1α/β^-/-^ corneas had less neutrophil infiltration than USA300 infected C57BL/6 corneas (**Fig 2B**). Further, USA300 infected IL-1α/β^-/-^ corneas show distinct bacterial aggregates in the corneal stroma (**Fig 2B,C**), indicating that the punctate keratitis observed in infected corneas of IL-1α/β^-/-^ mice is a result of these bacterial aggregates rather than infiltrating immune cells. Of the relatively few cells in infected IL-1α/β^-/-^ mice, infiltrating neutrophils did not appear to overlap with the dense aggregates in infected IL-1α/β^-/-^ corneas (**Fig 2C**). When compared with USA300 infected C57BL/6 mice, corneal thickness quantified microscopically was significantly less in ΔHla infected C57BL/6 and USA300 infected IL-1α/β^-/-^corneas (**Fig 2D**).

**Figure 2.**
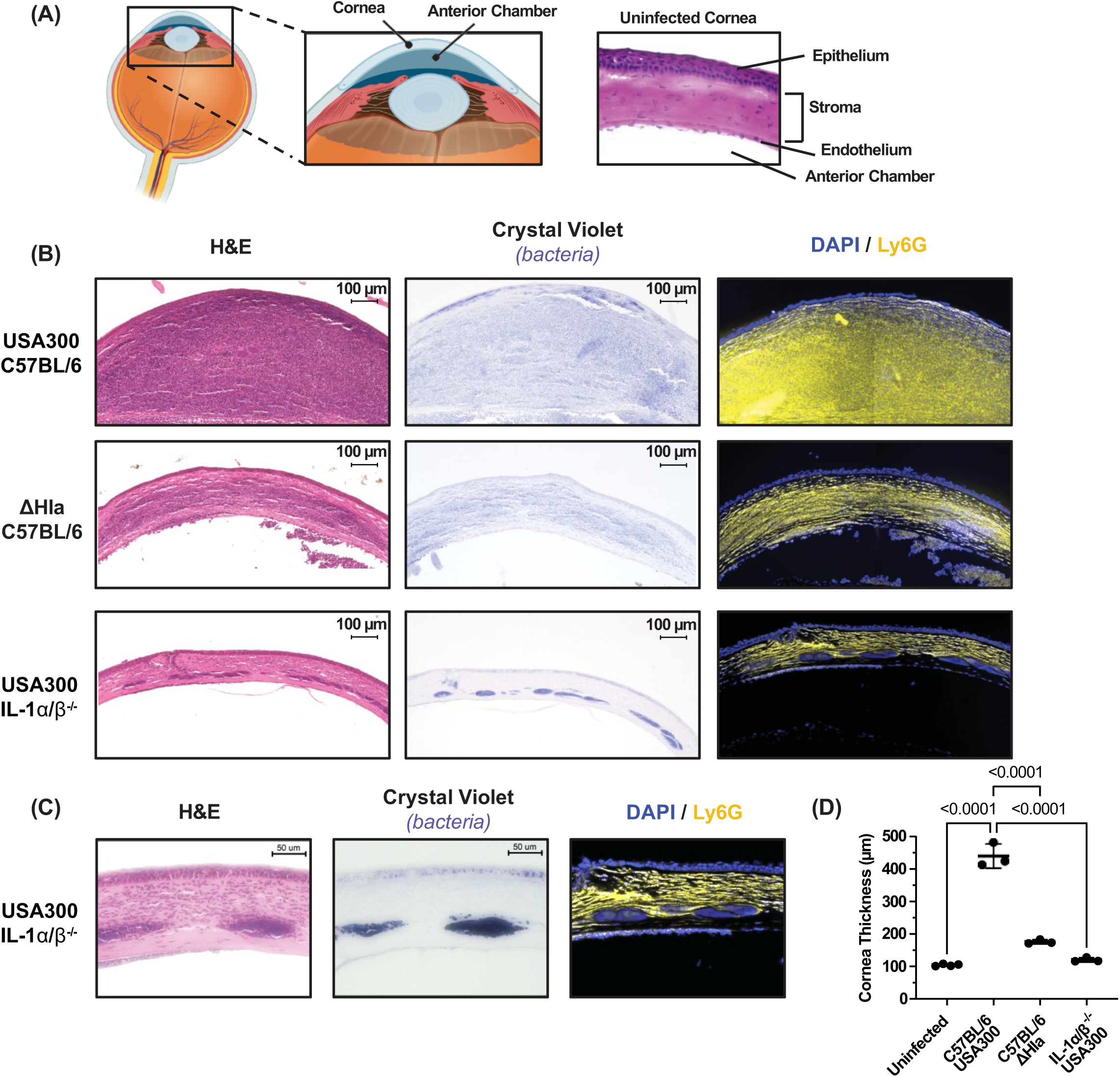
IL-1β and α-hemolysin regulate corneal thickness and neutrophil infiltration in MRSA infected corneas. Corneas of C57BL/6 or IL-1α/β^-/-^ were infected with 1x10^3^ CFU of USA300 or ΔHla. After 24 h eyes were sectioned, and stained by H&E or crystal violet to detect bacteria, or immunostained to identify Ly6G+ neutrophils. **A.** Diagram of the eye showcasing the cornea and a representative H&E image of an uninfected cornea. **B.** Representative low magnification (original 10x) images of H&E. **C.** Higher magnification (original 20x) representative images of USA300 infected IL-1α/β^-/-^ corneas highlighting bacterial aggregates in the corneal stroma. **D.** Quantification of corneal thickness by image analysis. Statistical significance was determined by 1-way ANOVA followed by Dunnett’s multiple comparison test.

Corneas infected with *S. aureus* strain 8325-4 generated a similar phenotype as USA300 in IL-1α/β^-/-^ compared with C57BL/6 mice. Punctate opacity, bacterial aggregates in the stroma, few infiltrating neutrophils, and increased CFU were all observed in the IL-1α/β^-/-^ corneas infected with 8325-4 (**Fig S1A-D**). Similarly, bacterial aggregates were detected in immune compromised IL-1R1^-/-^ and MyD88^-/-^ corneas (MyD88 is engaged following IL-1R activation) following epithelial abrasion and topical infection with 8325-4 (**Figure S1E)**, indicating that these bacterial aggregates occur following *S. aureus* infection in the absence of IL-1β signaling and are independent of the method of infection.

### Neutrophils are the major source of IL-1β in MRSA infected corneas

To quantify the number of infiltrating neutrophils and monocytes following infection with USA300 or ΔHla, corneas were dissected 24 h post infection and digested with collagenase. Total cells were examined by flow cytometry to identify CD45^+^myeloid cells, neutrophils (CD11b^+^Ly6G^+^), and monocytes (CD11b^+^Ly6C^+^). The gating strategy to derive live, single CD45^+^ cells is shown in **Fig S2,** and representative scatter plots are shown in **Fig 3A**. CD45^+^ cells comprised >90% total cells recovered from infected corneas, of which ∼90% were neutrophils and <10% were Ly6C^+^ monocytes (**Fig 3A,B**). C57BL/6 corneas infected with ΔHla and IL-1α/β^-/-^ corneas infected with USA300 had significantly less neutrophils than C57BL/6 corneas infected with USA300 (**Fig 3C**).

**Figure 3.**
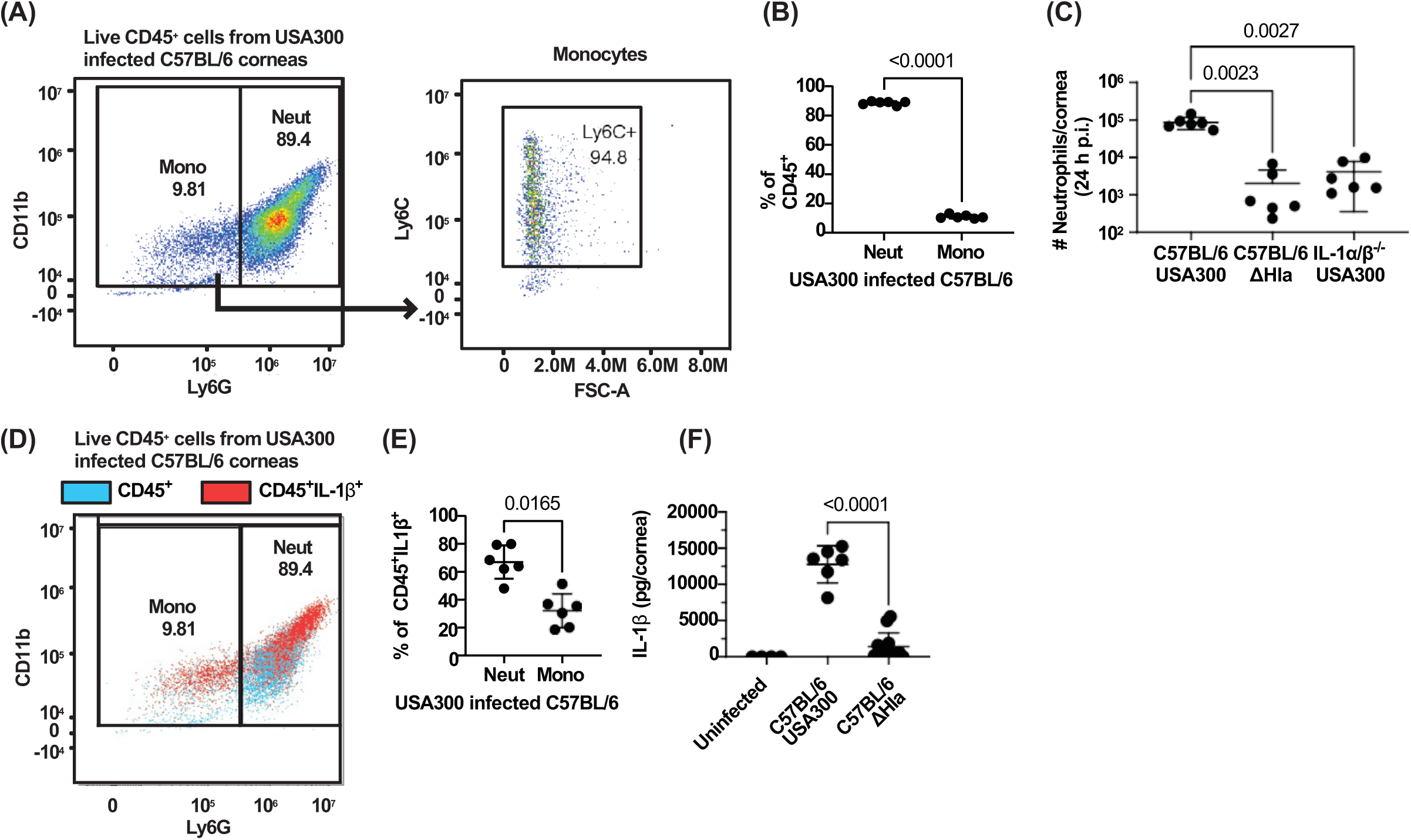
IL-1β and α-hemolysin regulate neutrophil recruitment to MRSA infected corneas. Cells recovered from infected corneas following collagenase digestion were processed for flow cytometry. **A**. Representative scatter plot showing neutrophils (CD45^+^CD11b^+^Ly6G^+^) and monocytes (CD45^+^CD11b^+^Ly6G^-^Ly6C^+^). **B.** Percent quantification of neutrophils and monocytes out of total CD45^+^ cells. **C.** Total number of neutrophils in USA300 and ΔHla infected C57BL/6 corneas and USA300 infected IL-1α/β^-/-^ corneas. **D,E:** Representative scatter plot and quantification of intracellular IL-1β in monocytes and neutrophils. **F.** IL-1β in infected corneas quantified by ELISA. Statistical significance was determined by 1-way ANOVA followed by Dunnett’s multiple comparison test **(C)**, paired T-test **(B,E)**, or unpaired T-test **(F)**. Error bars represent mean +/- SD. Data points represent individual corneas.

To identify the source of IL-1β in infected corneas, CD45^+^ cells from USA300 infected C57BL/6 corneas were also fixed and stained for intracellular IL-1β. Although both neutrophils and monocytes had IL-1β+ populations, neutrophils comprised the highest percent of IL-1β expressing cells **(Fig 3D,E)**. Infected corneas were also homogenized and IL-1β was quantified by ELISA. We found that infection with ΔHla induced significantly less IL-1β than infection with USA300 **(Fig 3F)**, likely because there were fewer neutrophils and monocytes.

Overall, these results highlight the important role for α-hemolysin in inducing neutrophil IL-1β secretion in infected corneas and that IL-1β regulates bacterial growth. However, in contrast to *P. aeruginosa* and *S. pneumoniae* keratitis^26,27^, the absence of IL-1β in *S. aureus* keratitis leads to the formation of distinct bacterial aggregates and minimal immune cell infiltrate in the corneal stroma, both of which are hallmarks of infectious crystalline keratopathy seen in corticosteroid treated patients^28–32^.

### α-hemolysin-induced IL-1β secretion by neutrophils requires NLRP3, caspase-1 and Gasdermin D (GSDMD)

Considering that neutrophils are the primary source of IL-1β during MRSA corneal infections, we investigated if α-hemolysin induces neutrophil IL-1β secretion under defined *in vitro* conditions and if this is dependent on the canonical NLRP3 pathway. α-hemolysin is secreted during late log and stationary phase of growth following activation of the quorum sensing accessory gene regulator (AGR) system ^33^. To detect the optimal time point for Hla release during bacterial growth, media were collected from USA300 and ΔHla cultures after 1.5 h, 4 h, and overnight. Cultures were centrifuged to remove bacteria, and supernatants were passed through 0.2 µm pore filters as described by Nunez et al . The resulting conditioned media was then precipitated by trichloroacetic acid (TCA) and examined by western blot using well characterized antibodies to α-hemolysin.

We found no differences in growth rate between USA300 and ΔHla (**Fig 4A)**, and Hla was not detected during early logarithmic growth (OD660 at 1.5h); however, the 33kD Hla monomer was detected after 4 h of growth and was highest after overnight incubation (**Fig 4B**). We also confirmed that α-hemolysin is absent in the ΔHla mutant strain. Subsequent *in vitro* experiments were performed using conditioned media from overnight cultures.

**Figure 4.**
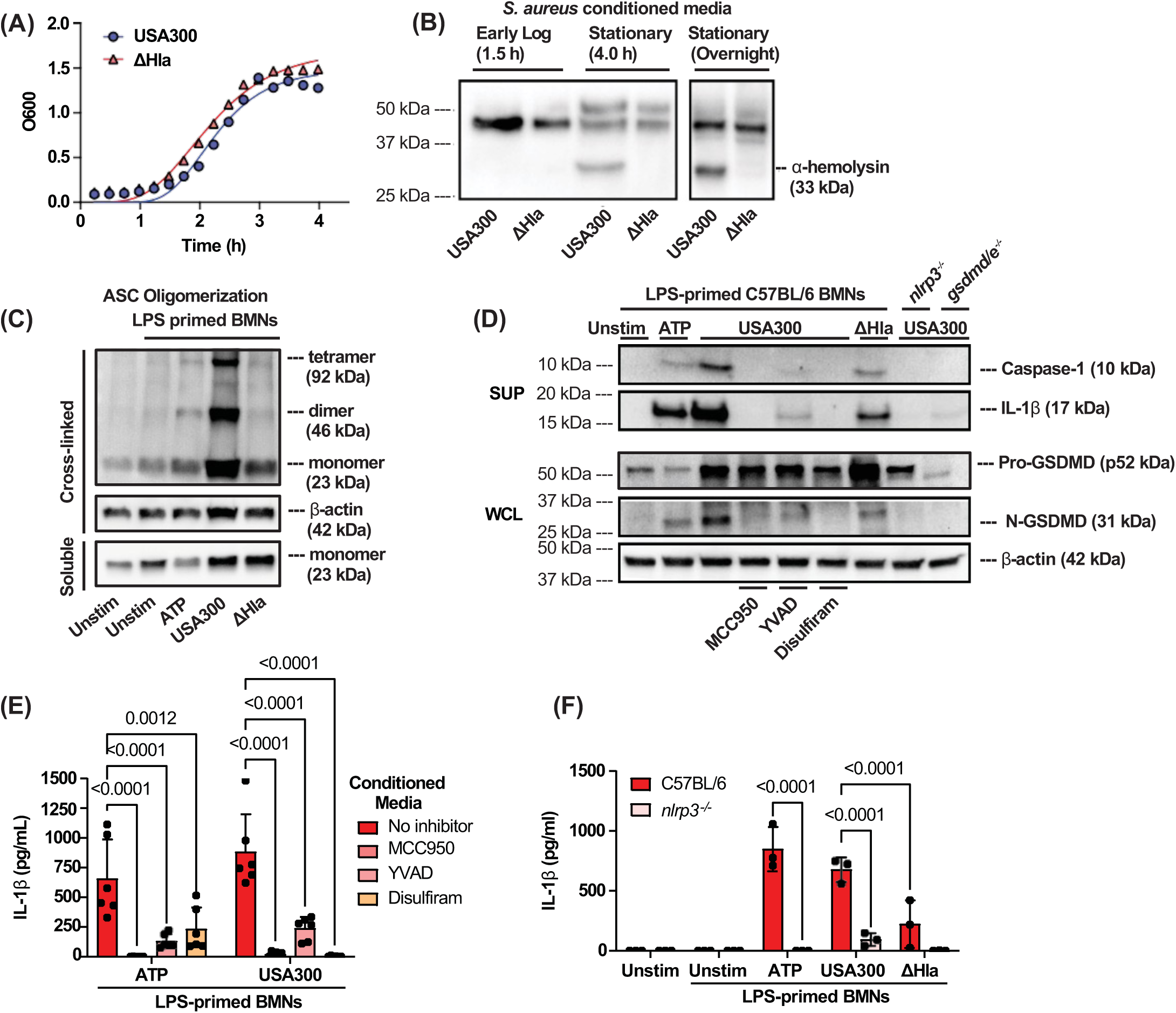
α-hemolysin (Hla) activates the NLRP3 inflammasome in murine neutrophils. **A.** USA300 and ΔHla growth curves (OD600) showing no difference between strains. **B. U**SA300 and ΔHla cultures were centrifuged and filtered to acquire cell-free conditioned media. α-hemolysin in conditioned media was assessed by western blotting. **C-F.** C57BL/6 or *nlrp3^-/-^*bone marrow neutrophils (BMNs) were primed for 3 h with 500 ng/mL LPS then stimulated for 1 h with 3 mM ATP or 1:10 diluted conditioned media from overnight cultures of USA300 or ΔHla. **C.** ASC oligomerization following cross-linking; ASC monomers shown in cross-linked and soluble fractions. **D.** Proteolytic cleavage of caspase-1, GSDMD, and IL-1β following stimulation in the presence of 2 µM NLRP3 inhibitor MCC950, 5 µM caspase-1 inhibitor YVAD, or 10 µM GSDMD inhibitor Disulfiram. **E-F.** IL-1β secretion quantified by ELISA following stimulation of C57BL/6 or *nlrp3^-/-^* BMNs or incubation in the presence of inhibitors. Statistical significance was determined by 2-way ANOVA with Šídák’s multiple comparison test **(E-F)**. Error bars represent mean +/- SD. Each data point represents biological replicates. Western blots represent at least two biological replicates.

The canonical NLRP3 pathway in macrophages and neutrophils requires ASC oligomerization, caspase-1 cleavage, and GSDMD cleavage; however, in contrast to macrophages, this pathway does not lead to pyroptotic cell death in LPS primed neutrophils stimulated with canonical NLRP3 activators ATP or nigericin^34^. To assess the role of α-hemolysin in ASC oligomerization, LPS-primed bone marrow neutrophils were examined after 1 h stimulation with conditioned media from overnight cultures of USA300 or ΔHla.

Oligomerization was detected by western blot analysis following cross-linking of ASC proteins. We found that while the ASC monomer was detected in soluble and cross-linked proteins in all conditions, the ASC dimer was only detected in neutrophils stimulated with ATP or with conditioned media from USA300, but not ΔHla cultures (**Fig 4C**).

To determine if conditioned media containing Hla induces cleavage of inflammasome associated proteins, bone marrow neutrophils from C57BL/6, *nlrp3*^-/-^, or *gsdmd/e^-/-^* mice were primed 3h with LPS and incubated for 1 h with conditioned media from USA300 or the ΔHla mutant. Neutrophils were also incubated with USA300 conditioned media in the presence of small molecule inhibitors of NLRP3 (MCC950), caspase-1 (YVAD), or GSDMD (disulfiram). Proteins from neutrophil culture supernatants were precipitated using TCA, and cleaved caspase-1 (p10) and IL-1β (p17) were detected by western blot analysis. Pro-GSDMD processing to N-GSDMD (31kD) in neutrophil lysates was also examined.

Cleaved IL-1β, caspase-1, and GSDMD were detected in neutrophils stimulated with ATP or with conditioned media from USA300, but not in the presence of MCC950, YVAD or disulfiram or in neutrophils from *nlrp3*^-/-^ or *gsdmd/e*^-/-^ mice (**Fig 4D**). Cleaved proteins were also detected in neutrophils stimulated with conditioned media from ΔHla mutants, but at lower levels (**Fig 4D**).

As a second approach to determine the role of α-hemolysin and the NLRP3 inflammasome in neutrophil IL-1β secretion, LPS-primed bone marrow neutrophils from C57BL/6 mice were incubated in the presence of these inhibitors and IL-1β secretion was quantified by ELISA using monoclonal antibodies that primarily detect cleaved IL-1β.

We found that USA300 conditioned media induced significantly less neutrophil IL-1β secretion in the presence of each of these inhibitors (**Fig 4E**). Similarly, USA300 conditioned media induced high levels of IL-1β secretion by C57BL/6 neutrophils that were completely absent in *nlrp3^-/-^* neutrophils (**Fig 4F**). Neutrophils treated with ΔHla conditioned media secreted significantly less IL-1β than those incubated with USA300 conditioned media, and was also absent in *nlrp3^-/-^*neutrophils (**Fig 4F**). We could not detect the pro-form of caspase-1 (45 kDa) or IL-1β (31 kDa) likely because of non-specific Fc binding by *S. aureus* Protein A (**Fig S6A,B**).

Collectively, these results demonstrate that α-hemolysin-induced neutrophil IL-1β secretion is dependent on activation of the canonical NLRP3 pathway. While α-hemolysin is necessary to induce robust NLRP3 pathway activation, lower levels of caspase-1, GSDMD, and IL-1β processing were visible in neutrophils treated with conditioned media from the ΔHla mutant, suggesting that other toxins can contribute to neutrophil NLRP3 activation. Indeed, an earlier report showed that while α-hemolysin is the primary inducer of IL-1β secretion by bone marrow-derived macrophages, deletion of α-, β-, and γ-hemolysins was necessary for complete ablation of IL-1β secretion^35^.

### Hemolysis and neutrophil IL-1β secretion induced by MRSA CC5 and CC8 clinical isolates is dependent on α-hemolysin

Gilmore, Andre and colleagues reported that MRSA clinical isolates from Mass Eye and Ear in their studies were mostly comprised of two clonal complexes CC5 from infected corneas, and CC8 from periocular skin. Genomic analysis of five CC5 and five CC8 clinical isolates from these studies demonstrated that all CC5 and CC8 isolates have a complete *Hla* gene; however, while all CC8 isolates have the same Hla sequence as the USA300 reference strain, all CC5 isolates used in this study harbor single nucleotide polymorphisms (SNPs) in the Hla gene and the TèC SNP at base 902 results in amino acid substitutions of residue 275 from isoleucine to threonine (I275T), whereas the TèG SNP at base 702 results in substitution of residue 208 from aspartic acid to glutamic acid (D208E) (**Table 1**, DNA sequence in **Fig S4**).

**Table 1.**
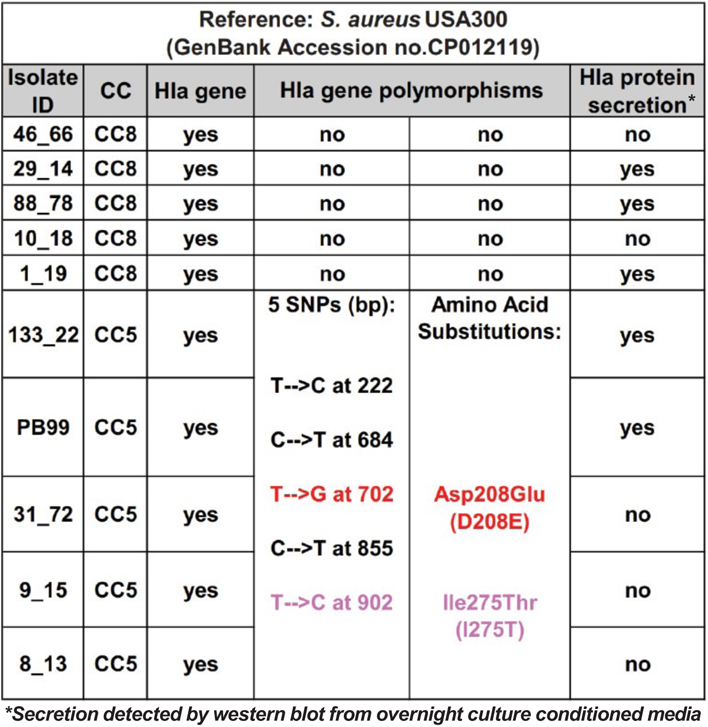
α-hemolysin polymorphisms in MRSA clinical isolates.

To examine Hla protein production in five CC5 and five CC8 of these clinical isolates, we generated conditioned media from overnight culture detected Hla by western blot. We found Hla in conditioned media from USA300 (CC8 strain) and in three of CC8 (although weakly in 1_19) and two CC5 clinical isolates (**Fig 5A)**. Hla was also detected in the lysates of isolates that secreted Hla (data not shown).

**Figure 5.**
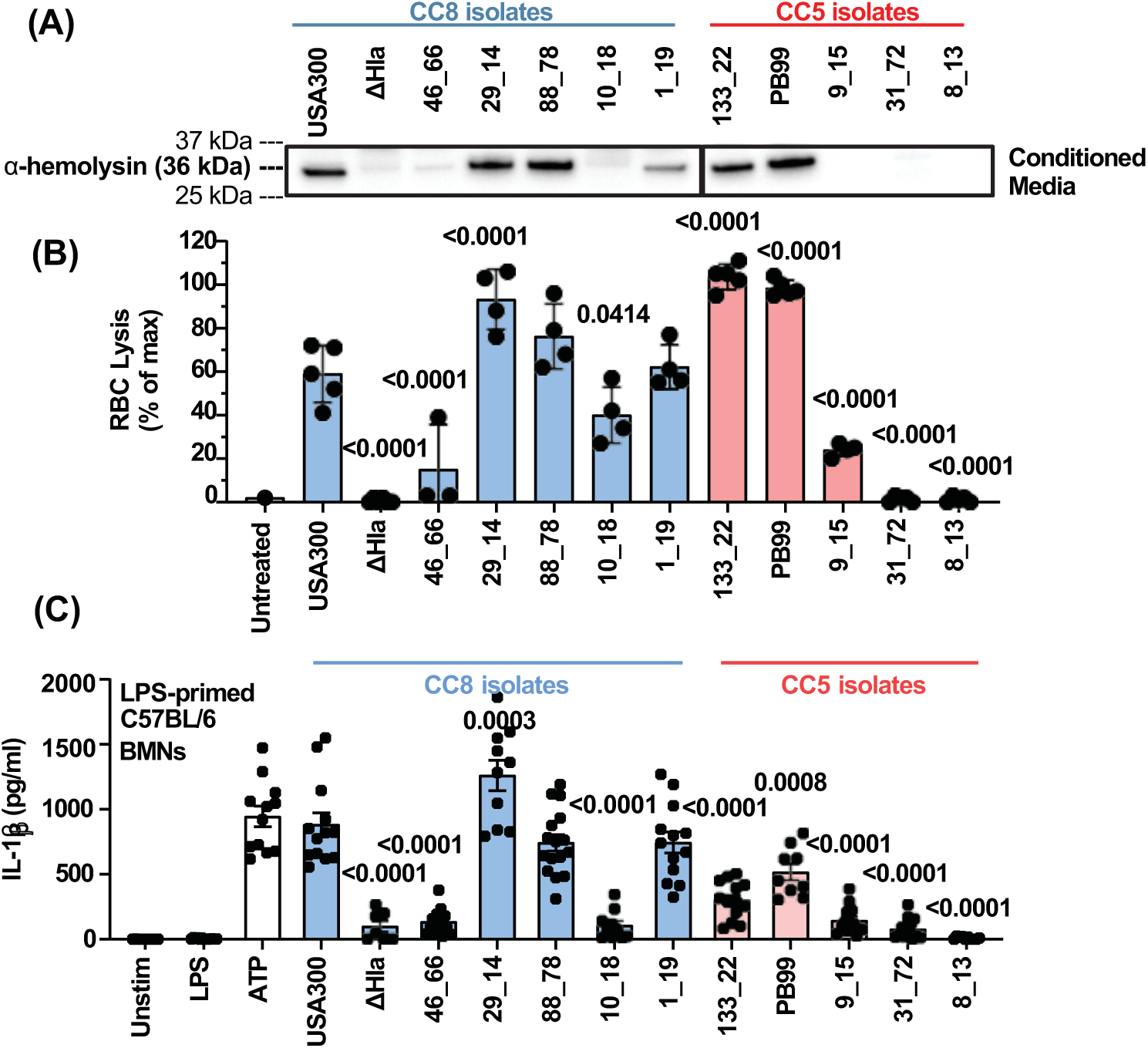
Rabbit RBC lysis and neutrophil IL-1β secretion is associated with α-hemolysin (Hla) secretion by MRSA clinical isolates. **A.** Conditioned media from overnight cultures of USA300, ΔHla, and CC5 and CC8 MRSA clinical isolates were precipitated by TCA and α-hemolysin was detected by western blot. **B.** Conditioned media were diluted 1:80 and incubated for 1 h with 10% rabbit RBCs. Following centrifugation, hemolysis was assessed by optical density. Results are percent of triton-x treated RBCs. **C.** IL-1β secretion by LPS-primed neutrophils from C57BL/6 mice following 1 h stimulation with 3 mM ATP or 1:10 diluted conditioned media from USA300, ΔHla, or MRSA clinical isolates. **B-C.** Statistical significance was determined by 1-way ANOVA with Dunnett’s multiple comparisons test using USA300 as the control comparison. Error bars represent mean +/- SD. Each data point represents biological replicates. Western blots represent at least two biological replicates.

Rabbit erythrocytes express high cell surface levels of the Hla receptor ADAM10 and are highly susceptible to lysis by α-hemolysin^16^. Conditioned media from USA300 (CC8), ΔHla and from CC5 and CC8 clinical isolates were incubated for 1 h with rabbit erythrocytes, and lysis was quantified by optical density at 570 nm. We found that Hla producing (Hla^+^) CC5 and CC8 isolates induce more hemolysis than non-Hla secreting (Hla^-^) isolates, with the extent of lysis approximately related to the amount of Hla in each isolate (**Fig 5B**). Hemolysis is also detected at high concentrations of conditioned media (**Fig S3A)**, possibly due to β- or γ-hemolysin.

To determine if Hla secreted by clinical isolates also regulates neutrophil IL-1β secretion, conditioned media from CC8 and CC5 clinical isolates, including USA300 and ΔHla strains, were incubated 1h with LPS-primed bone marrow neutrophils and IL-1β was quantified by ELISA. We found that IL-1β secretion was only induced by clinical isolates that secreted α-hemolysin (**Fig 5C**). IL-1β secretion induced by Hla^+^ clinical isolates was also inhibited in the presence of MCC950, YVAD, or disulfiram (**Fig S3B**), indicating that IL-1β secretion by Hla+ clinical isolates is also dependent on activation of the NLRP3 inflammasome.

We also show that conditioned media from both Hla^+^ and Hla^-^ clinical isolates stimulated release of similar levels of neutrophil elastase and myeloperoxidase (MPO) (**Fig S3B,C**), indicating that the release of these granule proteins is independent of Hla production. This is likely because mobilization of neutrophil granules is instead dependent on secretion of *S. aureus* phenol-soluble modulin (PSM) and staphylococcal superantigen-like (SSL) proteins that trigger neutrophil degranulation through activation of formyl peptide receptors^36–38^.

Collectively, these results demonstrate that α-hemolysin is the primary cause of RBC lysis and inducer of neutrophil IL-1β secretion by both CC5 and CC8 isolates.

### Hla determines corneal disease severity and bacterial growth in CC8, but not in CC5 infected corneas

To assess if α-hemolysin also determines corneal disease severity caused by CC5 and CC8 MRSA clinical isolates, corneas of C57BL/6 mice were infected with 1x10^3^ bacteria from Hla^+^ or Hla^-^ CC5 and CC8 isolates. Representative brightfield images show that Hla^+^ CC8 isolates 29_14 and 88_78 induce a localized intense corneal opacification that was comparable to USA300, whereas Hla^-^ CC8 isolates 10_18 and 46_66 had less and more diffuse corneal opacification, comparable to infection with ΔHla **(Fig 6A)**. 1_19 had an intermediate phenotype reflective of the lower Hla secretion shown by western blot in Figure 5. Similarly in CC5 clinical isolates, Hla^+^ PB99 caused severe disease and Hla- strains 31_72 and 8_13 caused mild disease; however, the correlation of Hla production and disease severity was not supported by Hla^+^ 133_22 which caused mild disease comparable to the ΔHla mutant, or by Hla- 9_15 which exhibited the more severe disease phenotype that is comparable to infection with USA300 (**Fig 6A**).

**Figure 6.**
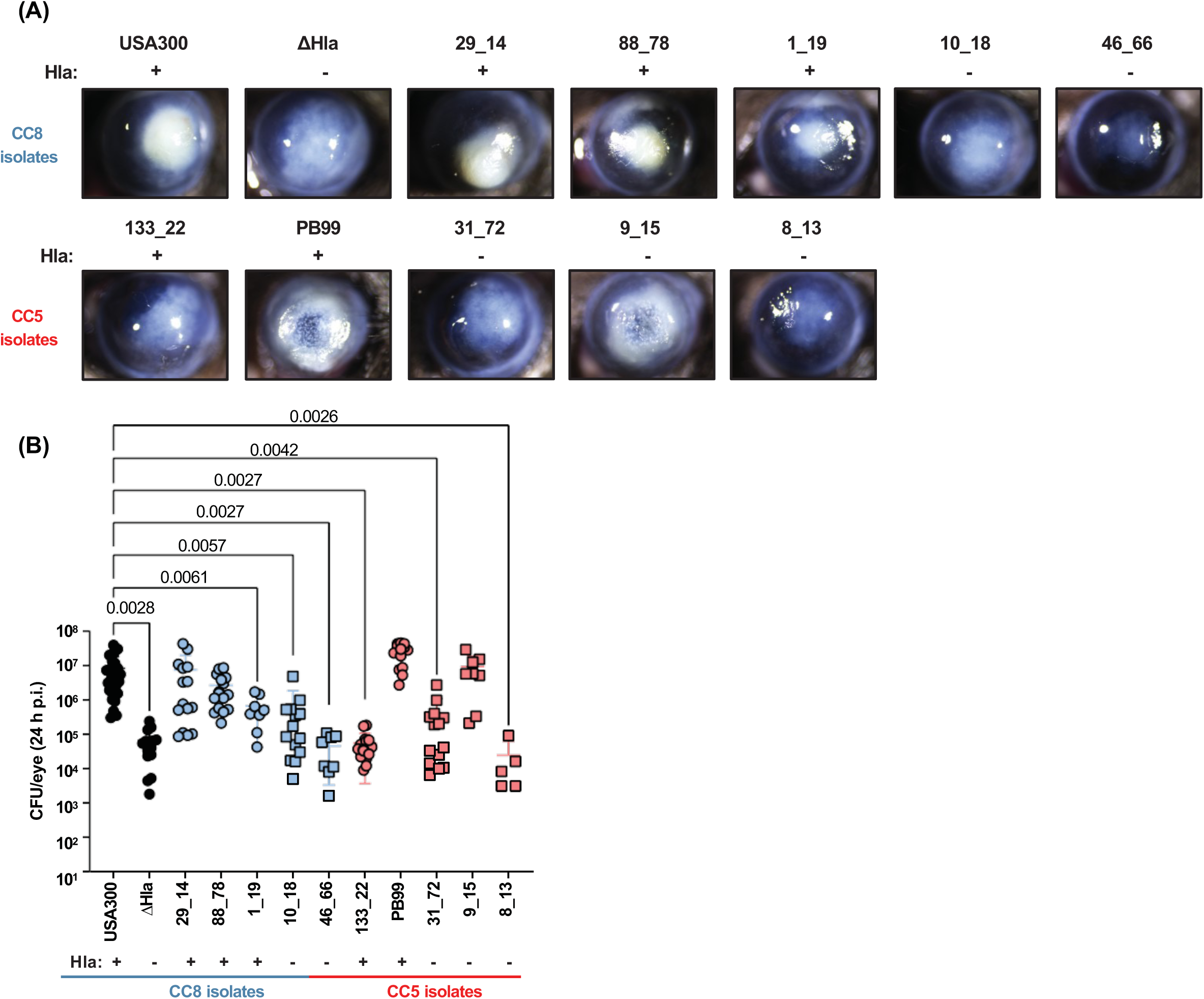
Corneal disease severity induced by MRSA CC8 isolates is associated with Hla. A 2 µL suspension of 1x10^3^ CFU USA300, ΔHla, or MRSA clinical isolates were injected into the corneal stroma of C57BL/6 mice. After 24 h, corneal disease and viable bacteria in infected corneas were assessed. **A.** Representative images of infected corneas. **B.** Quantification of viable bacteria in infected corneas by CFU. Statistically significant difference from USA300 was determined by 1-way ANOVA with Dunnett’s multiple comparisons test. Error bars represent mean +/- SD. Data points represent individual infected corneas.

To quantify viable bacteria following corneal infection with these clinical isolates, infected eyes were homogenized and CFU were counted and compared with corneas infected with USA300. As shown in **Fig 6B**, the bacterial load of Hla^+^ CC8 infected corneas was similar to USA300, whereas Hla- CC8 isolates were similar to the ΔHla mutant that caused milder disease and yielded significantly fewer CFU than USA300. Hla^+^ CC8 isolate 1_19 had an intermediate phenotype. In contrast to CC8 isolates, CFU from corneas infected with CC5 isolates were not directly related to Hla production as some Hla^+^ isolates such as 133-22 caused mild disease with less CFU and Hla^-^ isolates such as 9_15 caused severe disease.

These findings demonstrate an association between Hla expression *in vitro* and infection severity in CC8 infected corneas, despite the complexities of an *in vivo* infection. This contrasts with CC5 clinical isolates where corneal disease severity and bacterial growth were not closely associated with Hla secretion, indicating that either Hla is expressed during infection as they have the Hla gene, or that virulence factors other than Hla may be driving corneal disease severity caused by CC5 isolates.

### I275T and D208E substitutions in CC5 Hla protomers predict spontaneous oligomerization and altered ADAM10 binding

As noted above, all CC5 isolates used in this study harbor single nucleotide polymorphisms (SNPs) in the Hla gene that result in I275T and D208E amino acid substitutions (**Table 1, Fig S4**). Based on the published 3D structure of the Hla protomer, 3ANZ.pdb ^39^, we examined the predicted effect of the CC5 mutations on the protomer and the heptameric porin.

The predicted 3D structure of the N-terminal processed α-hemolysin protomer shows a predominantly beta-sandwich fold with compact globular cap and rim domains with an extended β-hairpin stem domain (**Fig 7A**). I275T and D208E substitutions are in the Hla cap and rim domains, respectively. The oligomerized α-hemolysin heptamer shows that these substitutions are radially distributed on the outside of the structure (**Fig 7B**). The D208E mutation is located in the membrane-interfacing rim domain of α-hemolysin (**Fig 7A,B)**, which is a structurally critical region that mediates plasma membrane binding and protomer oligomerization ^40^.

**Figure 7:**
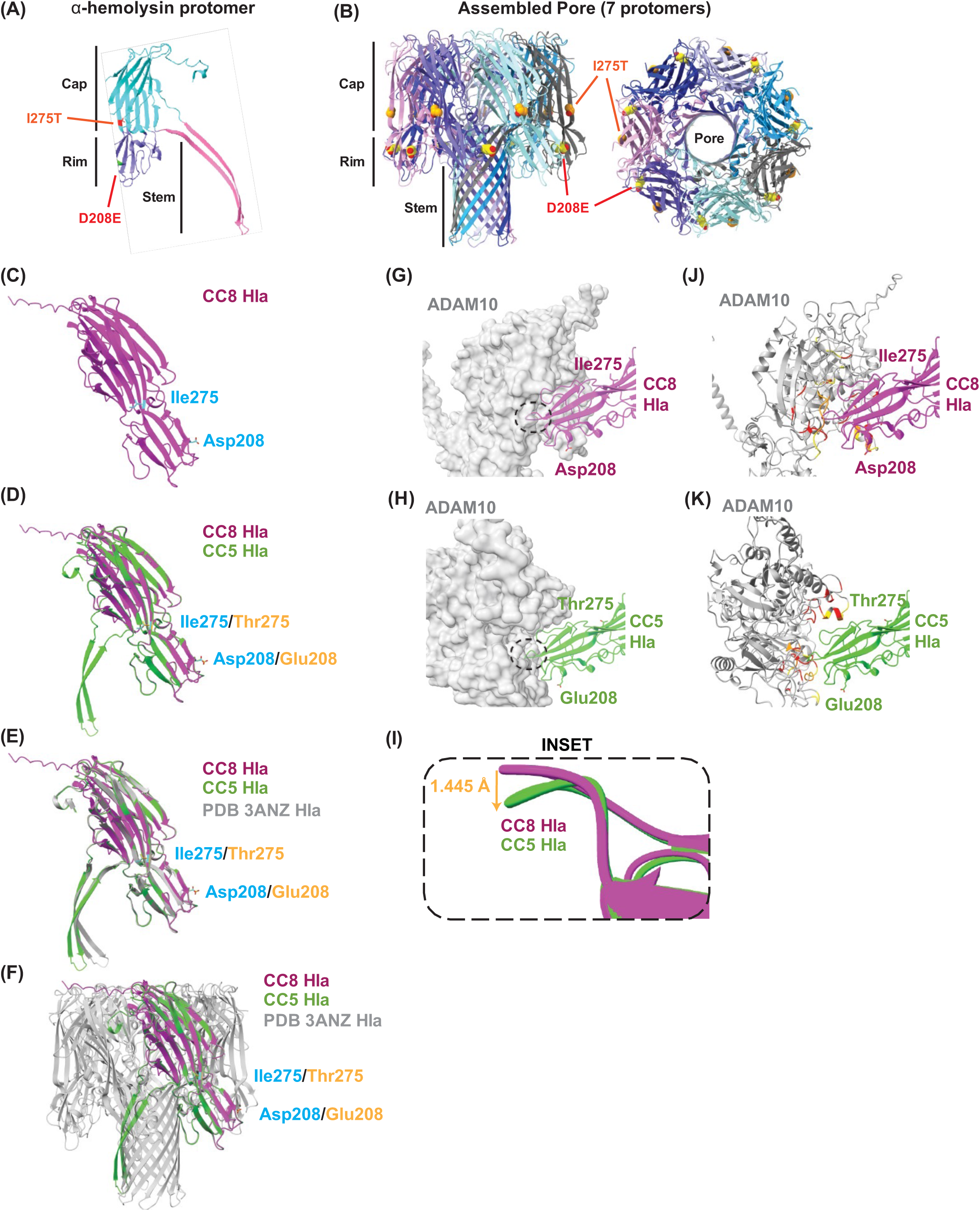
Predicted outcome of α-hemolysin mutations D208E/ I275T in CC5 clinical isolates. **A.** Schematic of α-hemolysin (Hla) protomer (PDB 3ANZ, Chain A) color-coded by the cap (cyan), rim (purple), and stem (pink) showing variants at positions 208 and 275. **B.** Schematic of heptameric Hla pore (PDB 3ANZ) color-coded by chain shown from the side (left) and top (right) with variants at positions 208 and 275 shown as spheres. **C.** AlphaFold3 model of CC8 Hla monomer (purple) showing isoleucine (Ile) at position 275 and aspartic acid (Asp) at position 208. **D.** AlphaFold3 model of CC5 Hla (green) overlaid on CC8 α-hemolysin (purple) showing CC5 variant Thr275 and Glu208 (orange) and CC8 variant Ile275 and Asp208 (cyan). **E.** CC8 and CC5 Hla overlaid with the published Hla monomer (grey, PDB 3ANZ). **F.** AlphaFold3 model of the full heptameric CC5 Hla pore (green) and the published Hla pore (grey). Variants at positions 208 and 275 are in orange and cyan as indicated. **G,H.** AlphaFold3 models of CC8 and CC5 Hla bound to ADAM10 showing CC8 Hla (purple, **G)** and CC5 Hla (green, **H**) bound toADAM10 (grey). Note the ∼90-degree rotation of the ADAM10 protein in the angle of ADAM10 binding between CC5 and CC8 Hla (**Movie S2**). Predicted sites of Hla/ADAM10 interactions are indicated. **I.** Zoomed-in view of interactions showing overlay of the CC8 and CC5 terminal loop of the rim domain. **J,K.** Secondary structure of ADAM10 from panel (G) showing ADAM10 regions within 5 angstroms of CC8 Hla (red), CC5 Hla (yellow), and within 5A of both models. **J.** modeled binding to CC8 Hla; and **K.** model based on CC5 binding with ADAM10 at a different angle.

To predict the effect of these α-hemolysin amino acid substitutions D208E and I275T on protein structure and receptor binding affinity, we employed UCSF ChimeraX^41^ and AlphaFold3^42^ to analyze the conformation of the α-hemolysin monomer,. Simulations with the CC8 monomer show a well-folded monomer with the stem folded up near the cap (**Fig 7C**), which is an inactive conformation^44^. However, when the same structure prediction was run with the D208E/I275T substitutions seen in CC5 isolates, the stem domain flipped out and away from the cap (**Fig 7D**). Superposition of CC5 Hla containing both I275T and D208E variants showed that all 3 domains, including the stem region, superpose well with the published structure of the protomer of the α-hemolysin pore (3ANZ.pdb) ^39^. Superposition was highly consistent, as reflected by a low resulted in a root mean square deviation (RMSD) of 0.42 Å, indicating very similar structures (**Fig 7E, Fig S5, Movie S1**). We therefore predict that the CC5 Hla is structurally similar to an already active protomer in a fully oligomerized α-hemolysin pore (**Fig 7F**). The stem domain being flipped out in the D208E/I275T monomer suggests that these substitutions put Hla in a state that can spontaneously oligomerize.

Considering these conformational changes, we predicted that D208E/I275T substitutions also alter the interactions of Hla with the ADAM10 cell surface receptor that facilitates α-hemolysin interactions with the plasma membrane^16^. Simulations on CC8 (D208/ I275) and CC5 (D208E/ I275T) variants of the monomer bound to ADAM10 (sequence from Uniprot O14672) predict direct binding between α-hemolysin and ADAM10, with high confidence prediction scores for both variants of α-hemolysin monomers (**Fig S5**). The bottom of the rim region has a lower confidence prediction score, though that is expected given the known looping/disorder in that region. In both models, this rim region is also predicted to bind ADAM10. However, simulations with the D208E/I275T Hla predict a ∼90-degree rotation of the ADAM10 protein, indicating differences in the binding interface between ADAM10 and Hla in CC5 compared with CC8 variants (**Fig 7G-K, Movie S2**).

Evaluation of these models revealed that overall alignment was good, with most of the structure matching closely Evaluation of these models revealed very similar α-hemolysin structures, with a Root Mean Squared Deviation (RMSD) of ∼0.4 angstroms (low RMSD indicates close alignment), although a minor region at the rim had a higher RMSD of 3.2Å. There were marked differences in the bottom of the rim, which comes into direct contact with ADAM10 at slightly different binding interfaces. At the end of the rim region, the loop between residues 260 and 266 shifts 1.445 angstroms down in the CC5 Hla compared to the CC8 Hla (**Fig 7I**). Overall, these simulations predict that D208E/I275T substitutions lead to changes in the conformation of Hla and the ADAM10-Hla binding interface that may impact the ability of Hla to oligomerize either spontaneously or through altered ADAM10 binding.

### ADAM10 inhibition blocks neutrophil IL-1β secretion induced by Hla+ CC8 but not Hla+ CC5 MRSA clinical isolates

The ADAM10 Hla receptor is expressed by multiple cell types, and RNA-seq data from the Immunological Genome Project (IMMGEN) shows elevated gene expression in neutrophils isolated from the inflamed peritoneal cavity^43^. To determine if ADAM10 is present on the neutrophil surface, we used flow cytometry to quantify ADAM10 on bone marrow neutrophils and neutrophils isolated from the peritoneal cavity following induction of sterile inflammation. While ADAM10 was detected on bone marrow neutrophils (higher than FMO), cell surface expression was higher in neutrophils from the inflamed peritoneal cavity (**Fig 8A**).

**Figure 8.**
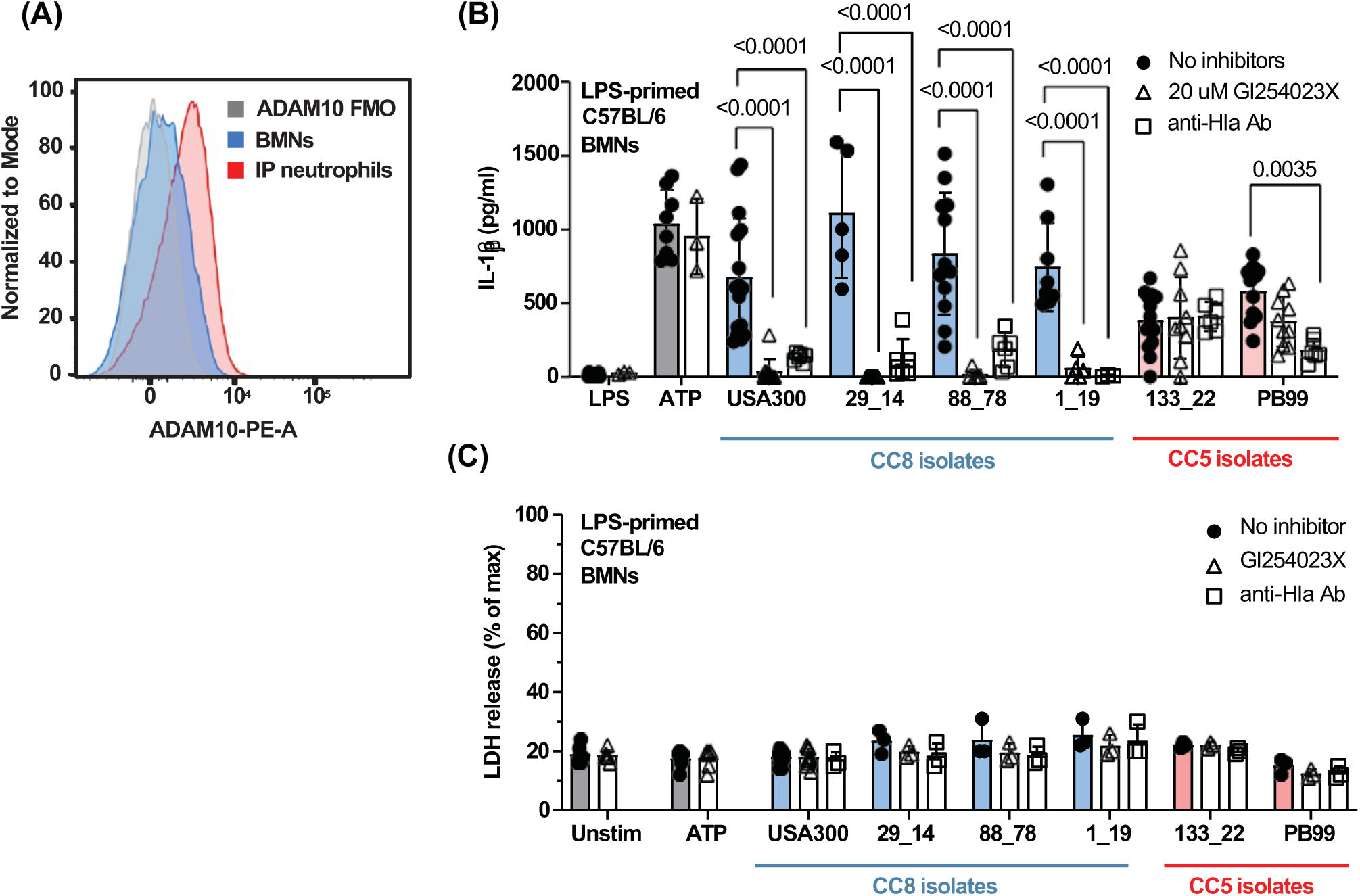
ADAM10 inhibition blocks neutrophil IL-1β secretion induced by Hla^+^ CC8 but not Hla^+^ CC5 isolates. **A.** Representative flow cytometry histogram showing expression of ADAM10 on unstimulated bone marrow neutrophils or peritoneal neutrophils following sterile inflammation. **B-C.** LPS-primed BMNs from C57BL/6 mice were stimulated for 1 h with conditioned media from USA300 or MRSA clinical isolates in the presence of 20 µM GI254023X (ADAM10 inhibitor) or with a neutralizing anti-Hla monoclonal antibody. IL-1β secretion and LDH release were quantified. Statistical significance was determined by 1-way ANOVA with Šídák’s multiple comparison test. Error bars represent mean +/- SD. Data points represent biological replicates.

To ascertain the role of ADAM10 on Hla-mediated neutrophil IL-1β secretion, we used LPS-primed bone marrow neutrophils as described for *in vitro* assays. Neutrophils were incubated 3 h with LPS and the small molecule ADAM10 inhibitor GI254023X ^44^. Neutrophils were then incubated 1 h with conditioned media from Hla^+^ CC5 or Hla^+^ CC8 clinical isolates and IL-1β secretion was quantified by ELISA. We found that IL-1β secretion by neutrophils stimulated with conditioned media from USA300 and Hla^+^ CC8 isolates was significantly inhibited in the presence of GI254023X; however, the inhibitor did not block IL-1β secretion by neutrophils stimulated with conditioned media from Hla^+^ CC5 isolates (**Fig 8B)**.

To confirm that these effects are specifically due to Hla, we incubated conditioned media with neutralizing monoclonal antibody to Hla prior to neutrophil stimulation. Anti-Hla antibody completely inhibited neutrophil IL-1β secretion induced by conditioned media from CC8 isolates, indicating that IL-1β secretion is specifically induced by Hla. Surprisingly, the anti-Hla antibody had no effect on conditioned media from 133_22 but significantly reduced IL-1β secretion induced by PB99 (**Fig 8B**). There were also no significant differences in neutrophil cell death induced by MRSA strains as measured by release of lactate dehydrogenase (LDH), which was ∼20% total (detergent treated neutrophils), and was not significantly different from unstimulated neutrophils (**Fig 8C**).

Collectively, results from these functional assays support predictions from structural analysis as the CC5 Hla variant was unaffected by ADAM10 inhibitors, which is either because spontaneous Hla oligomerization is ADAM10-independent or because changes in Hla-ADAM10 binding makes Hla resistant to ADAM10 inhibitors.

## DISCUSSION

Hla is a pore-forming toxin that causes efflux of ATP and potassium ^12,13^ and induces NLRP3-dependent IL-1β secretion by macrophages, leading to pyroptotic cell death^35,45,46^. In the current study, we demonstrate that Hla is important in neutrophil secretion of IL-1β following assembly of the NLRP3 inflammasome, and that neutrophil derived IL-1β plays an important role in a murine model of MRSA keratitis where it regulates corneal disease severity and bacterial killing. Although neutrophils are the predominant cell types in early stages of disease, most studies on inflammasome activity have focused on macrophages. However, we and others have identified striking differences in inflammasome activation between neutrophils and macrophages, including that whereas NLRP3 activation of macrophages results in pyroptotic cell death, neutrophils were the primary source of IL-1β following canonical NLRP3 activation or corneal infections with *P. aeruginosa, Streptococcus pneumoniae,* or *Salmonella typhimurium* and that neutrophils activated under identical *in vitro* conditions as macrophages do not undergo pyroptosis following canonical NLRP3 activation ^47–50^.

We subsequently used an N-GSDMD specific antibody to demonstrate that whereas N-GSDMD inserts into the macrophage plasma membrane, in neutrophils it localizes instead to azurophilic granules, and IL-1β secretion is dependent on autophagy proteins rather than pyroptotic cell death ^51^. More recently, we reported that in contrast to *Pseudomonas aeruginosa* mediated NLRC4 activation in macrophages, NLRP3 is activated in neutrophils due to the ADP ribosyl transferase activity of *P. aeruginosa* exotoxin S (ExoS).

In the current study, we demonstrate a critical role for neutrophil derived IL-1β and bacterial Hla in regulating the disease severity of MRSA keratitis. Although all MRSA clinical isolates used in this study have an intact *hla* gene, our results show that several CC8 and CC5 clinical isolates do not secrete α-hemolysin, which is consistent with prior studies showing differences in RNA expression among clinical isolates^20^. We found that clinical isolates that do not secrete α-hemolysin also induce less neutrophil IL-1β secretion, which is consistent with earlier studies showing similar results for human and murine monocytes^45^.

Considering the important contribution of α-hemolysin to disease severity in a murine model of keratitis, we hypothesized that CC8 and CC5 groups would have different patterns of α-hemolysin expression that correlate with their propensity to infect the cornea or periocular dermal tissues^24^. We found that corneal disease severity caused by CC8 clinical isolates is associated with Hla secretion as Hla^-^ strains caused less corneal disease than Hla^+^ strains.

However, the CC5 isolates, which were primarily isolated from infected patient corneas ^24^, showed little association between disease severity and Hla expression. This finding suggests that *in vitro* expression of Hla does not predict disease severity in CC5 clinical isolates. Overall, these results highlight the heterogeneity among MRSA lineages, similar to prior findings^19,52^.

Our findings also provide insight into potential binding of Hla to its ADAM10 receptor. ADAM10 is constitutively expressed on the surface of multiple cell types, including epithelial and vascular endothelial cells, where the protease activity plays an important role in ectodomain shedding by cleaving cell surface proteins such as ICAM-1^53,54^. ADAM10 activation by Hla facilitates pore formation in the plasma membrane^16,55^. Although we show here that while ADAM10 is expressed on neutrophils, Hla binding does not lead to cell death.

In all CC5 MRSA isolates examined in this study, we identified two α-hemolysin single nucleotide polymorphisms (SNPs) that result in amino acid substitutions I275T and D208E. Our structural modeling predicts that these substitutions cause the stem domain to flip out from the CC5 Hla monomer, causing the monomer to more readily adopt the active conformation seen in the assembled pore. This suggests that the CC5 Hla variant can form the active pore independent of ADAM10. In addition, conformational changes to CC5 Hla are predicted to alter the Hla-ADAM10 binding interface, causing ADAM10 to bind CC5 Hla and CC8 Hla differentially.

Functional analysis supports both models as neutrophil IL-1β secretion induced by CC8 Hla, but not the CC5 D208E/I275T Hla, was blocked by the ADAM10 inhibitor GI254023X. This finding can be explained either by spontaneous oligomerization of the D208E/I275T variant that does not require ADAM10, or that the changes in the ADAM10-Hla interface in the D208E/I275T variant impacts the ability of GI254023X to inhibit IL-1β secretion. Based on these findings, **Figure 9** predicted sequence of events shows that Hla from CC8 MRSA isolates binds ADAM10, which mediates Hla oligomerization and pore formation, resulting in activation of the NLRP3 inflammasome and IL-1β cleavage and release. In contrast, the CC5 Hla is not dependent on ADAM10, although it also induces IL-1β secretion through the canonical NLRP3 pathway.

**Figure 9.**
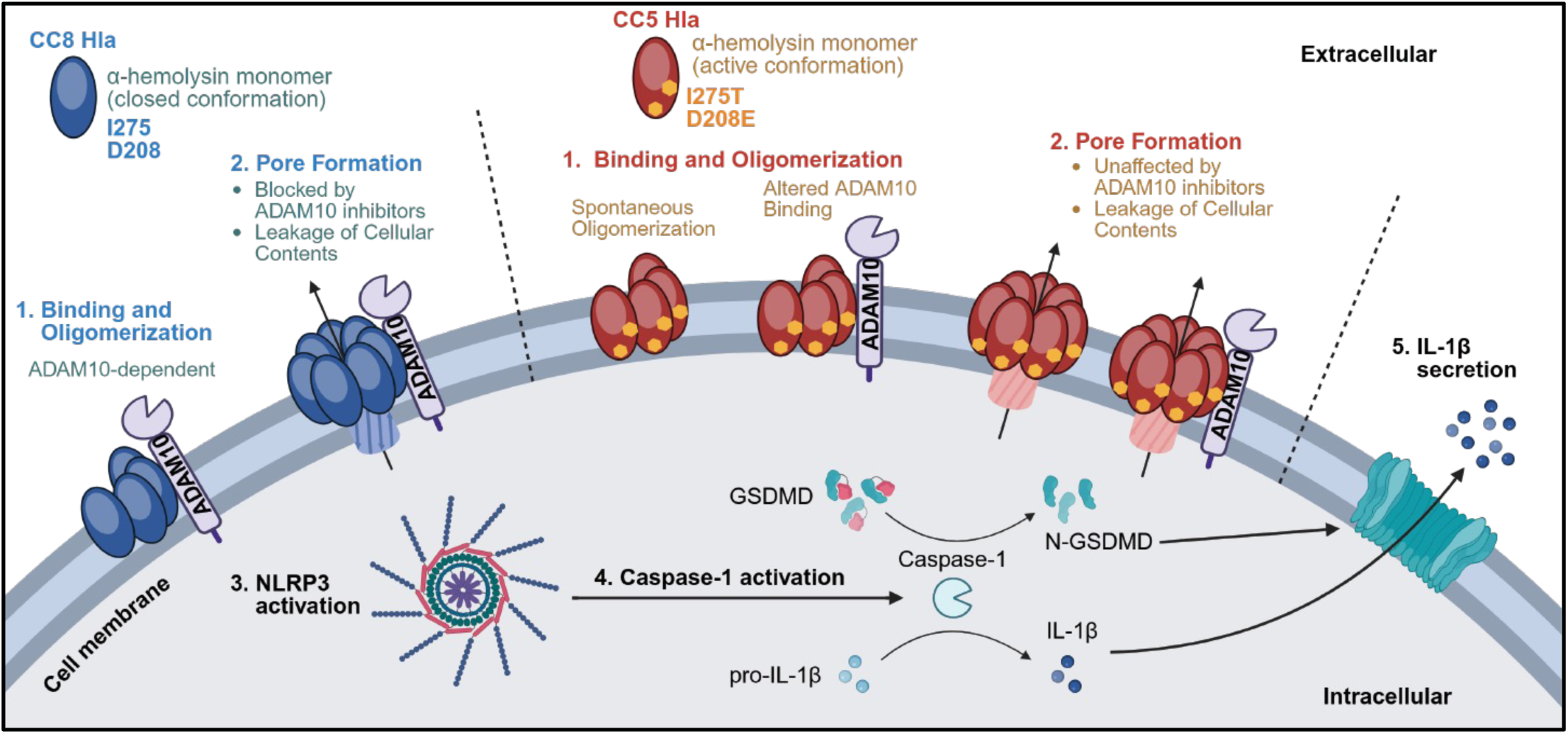
Predicted sequence of events. α-hemolysin from USA300 and CC8s are predicted to bind the ADAM10 receptor, transition into the active conformation, and oligomerize into the heptamer pore. Pore formation leads to leakage of intracellular contents which activates the NLRP3 inflammasome leading to subsequent IL-1β processing and secretion. In contrast, CC5 Hla with D208E/I275T substitutions is predicted to be in an already active conformation, potentially oligomerizing into the pore independent of ADAM10 or with altered ADAM10 binding. Both models are supported by functional data showing that neutrophil IL-1β secretion induced by CC5 Hla is unaffected by ADAM10 inhibitors. Created in BioRender. Liboro, K. (2026).

While several reports have identified residues important for α-hemolysin function, few have looked at these residues in the context of ADAM10 binding. For example, the amino acid substitutions identified in this report are strikingly close to residues Asp276 and Arg200 that are critical for α-hemolysin binding and which were identified prior to the discovery of ADAM10 as the receptor ^56,57^. Similarly, these findings support the results of a recent *in silico* study that predicted that residue 208 could mediate ADAM10-α-hemolysin binding ^58^. To the best of our knowledge, ours is also the first study showing ADAM10 cell surface expression in neutrophils and a functional role for ADAM10-α-hemolysin-mediated IL-1β secretion in neutrophils. Future studies will use site-directed mutagenesis and molecular binding studies to determine the impact of D208E/I275T in α-hemolysin-ADAM10 binding kinetics.

In conclusion, the results of this study identify essential roles for α-hemolysin and neutrophil IL-1β secretion in keratitis caused by MRSA. We also present predictive and functional insights into the molecular interactions between Hla variants and the ADAM10 receptor. α-hemolysin-ADAM10 binding interactions represent potential novel targets for Hla mediated intervention in *S. aureus* keratitis and other infections.

## MATERIALS AND METHODS

### Source of mice

C57BL/6 and *nlrp3^-/-^* mice were purchased from the Jackson Laboratories. *Gsdmd^-/-^gsdme^-/-^* mice were generated in-house by crossing *gsdme^-/-^* mice purchased from Jackson Laboratories and *gsdmd^-/-^* mice generously provided to us by Dr. Russel Vance (University of California, Berkeley). *Il1α^-/-^*, *Il1β^-/-^, Il1α^-/-^/β^-/-^,* and *il1-r1*^-/-^ mice were generated by Dr. Y. Iwakura (University of Tokyo). MyD88-deficient mice were generated by Shizuo Akira (Research Institute for Microbial Disease, Osaka University, Osaka, Japan).

All transgenic mice were on a C57BL/6 background. Male and female mice ages 6-12 weeks old were used for experiments. All animals were bred and housed according to institutional guidelines by the University of California, Irvine IACUC.

### Source of Methicillin resistant *S. aureus*

USA300 and MRSA clinical isolates (**Table S1**) were obtained from Mass Eye and Ear Hospital, Boston MA). The ΔHla mutant is NE1354 from the Nebraska collection generated by transposon insertion into *hla* ^59–61^.

### Isolation of murine bone marrow neutrophils (BMNs) and intraperitoneal (IP) neutrophils

For IP neutrophils, mice were injected intraperitoneally with two doses of 1 mL casein solution (9% w/v casein, 0.1375 mM MgCl2, 0.45 mM CaCl2, 1x PBS). The first dose was administered the day before and the second dose was administered 3 h prior to harvesting IP lavage. For BMNs, bone marrow from the tibias and femurs of C57BL/6 mice were collected. IP lavage and bone marrow were enriched for neutrophils using negative selection magnetic beads following the manufacturer’s instructions (STEMCELL, Milteny). These protocols yield >85% neutrophils as determined by flow cytometry

### *In vitro* stimulation conditions

USA300, ΔHla, or MRSA clinical isolate frozen stocks were streaked onto brain heart infusion agar (BHI; Millipore 53286, BD Difco 241830) and incubated at 37°C for 24 h to allow colony formation. Overnight cultures were made by inoculating 3 mL BHI broth with three picked colonies and incubating at 37°C at 200 rpm for 18-24 h. Conditioned media was prepared by centrifuging *S. aureus* overnight cultures twice at 3000 x g for 10 min to pellet the cells and filtering supernatants with a 0.22 µm pore filter (Genesee Scientific, Cat# 25-244). For Hla neutralization experiments, conditioned media was incubated with anti-α-hemolysin antibody (1:500; Abcam, ab190467) for 30 min prior to addition to neutrophils or rabbit RBCs.

After neutrophil enrichment, BMNs were resuspended in RPMI media containing 2% FBS and 20 ng/mL murine GM-CSF (StemCell, Cat# 78017.1) then incubated for 3 h with 500 ng/mL LPS (InvivoGen, 5974-44-02) to induce transcriptional expression of NLRP3 and pro-IL-1β. BMNs were then centrifuged and the media was replaced with RPMI without additives.

BMNs were incubated for 30 min with either 2 µM MCC950 (NLRP3 inhibitor; InvivoGen, Cat# inh-mcc), 5 µM YVAD (caspase-1 inhibitor; Sigma-Aldrich, SML0429-5MG), or 10 µM Disulfiram (GSDMD inhibitor; MedChem Express, HY-B0240/CS-2209). BMNs were then treated for 1 h with 3 mM ATP (canonical activator of NLRP3; Sigma-Aldrich, A6419-5G) or incubated with *S. aureus* conditioned media (1:10 final dilution). For experiments involving ADAM10 inhibition, neutrophils were incubated with 10 µM of GI254023X for 3.5 h (occurs simultaneously with LPS-priming) before stimulation with ATP or *S. aureus* conditioned media.

### Bacterial growth curves

USA300 and ΔHla colonies were picked and inoculated in 1 mL BHI media in a 12-well plate. The plate was incubated at 37 C with continuous shaking in a Cytation5 Imaging Reader (Biotek) with OD600 absorbance readings taken every 15 min. A nonlinear regression using a Gompertz growth curve was fitted onto the data.

### Quantification of Neutrophil Elastase, Myeloperoxidase, and IL-1β release by ELISA and cell death by LDH release

BMNs were plated in 96-well plates at 200,000 cells/well, primed with LPS, and stimulated according to *in vitro* stimulation procedures described above. Cell-free supernatants were collected by centrifuging at 300 x g for 10 min in V-bottom 96-well plates (PlateOne).

Neutrophil Elastase, Myeloperoxidase, and IL-1β release were measured by ELISA following manufacturer’s instructions (R&D DuoSet; DY4517, DY3667, DY401). Cell cytotoxicity was measured by LDH release assay following manufacturer’s instructions (Promega, G1781).

Percentage of cell death was determined based on maximum LDH release following 30 min incubation with cell lysis solution (Promega, G1821).

### Western Blot and ASC Oligomerization

BMNs were plated in 6-well plates at 2x10^6^ cells/well, primed with LPS, and stimulated according to *in vitro* stimulation procedures described above. Cell supernatants were collected, and protein was precipitated using 10% sodium deoxycholate and 100% trichloroacetic acid (TCA). The resulting pellets were washed with 100% acetone and solubilized in 0.2 M NaOH. Whole cell lysates (WCL) were collected by incubating cell pellets for 10 min on ice with 1x lysis buffer (Cell Signaling Technology, #9803) or 1x RIPA buffer (Sigma Aldrich, R0278-50mL) containing 5 mM diisopropylfluorophosphate (DFP), a potent inhibitor of serine proteases. The addition of DFP inhibits neutrophil proteases that may process target proteins such as caspase-1, GSDMD, and IL-1β after lysis. WCLs were then centrifuged at 16,000 rcf x g for 10 min at 4°C and clear supernatants were collected. Protein concentrations were determined by bicinchoninic acid (BCA) assay (Thermofisher, 23225). WCL were mixed with Laemmli buffer (Bio-Rad, #1610747) containing β-mercaptoethanol (2.5% final concentration) (Sigma-Aldrich, M3148).

For assessing ASC Oligomerization, WCL were instead centrifuged at 13,200 rpm for 20 min and the resulting clear supernatants were incubated at room temperature for 30 min with 4 mM disuccinimidyl suberate (DSS) to cross-link proteins. Cross-linked WCL is centrifuged again at 13,200 rpm for 20 min and the resulting pellet is resuspended in Laemmli buffer containing β-mercaptoethanol.

Unless otherwise noted, 20 µg of WCL, 25 µL of cross-linked WCL, or 20 µL of precipitated supernatants were separated on pre-cast 12% acrylamide gels (Bio-Rad, #4561043) via SDS-PAGE gel electrophoresis. Protein was transferred to PVDF membranes (Millipore, IPVH00010) using the TransBlot Turbo semi-dry apparatus (Bio-Rad). Western blot membranes were probed for caspase-1 (Abcam, ab179515), IL-1β (R&D, AF-401-NA), GSDMD (Abcam, ab209845), β-actin (Santa Cruz, sc-4777), or ASC (Adipogen, AG-25B-0006-C100). For detection of total protein, membranes were incubated for 5 min with Ponceau S solution (Sigma, P7170-1L) then washed with milliQ water before brightfield imaging on a ChemiDoc MP Imaging System (Bio-Rad).

For detection of α-hemolysin, protein in conditioned media were precipitated by TCA and then western blotted as described above. Western blot membranes were stained with ponceau S solution (Sigma Life Sciences, P7170-1L) for detection of total protein and probed using anti-α-hemolysin antibody (abcam, ab190467).

### Rabbit Hemolysis Assay

*S. aureus* conditioned media were incubated 30 min prior to serial dilution in PBS in a 96-well plate. Rabbit RBCs (Innovative Research, IRBRBC10ML) were centrifuged at 220 rpm for 10 min before aspirating the supernatant to remove free-floating hemoglobin and washed x3.

RBCs were added at a final concentration of 10% to the 96-well plates containing serially diluted conditioned media. Plates were incubated 1h at 37C before centrifuging the plate to pellet intact RBCs. Supernatants were transferred to a fresh 96-well plate and absorbance was measured at 570 nm. Percentage of RBC lysis was determined by normalizing absorbance against RBCs treated with 1x triton for 100% lysis.

### Murine model of *S. aureus* corneal infection

USA300, ΔHla, MRSA clinical isolates, or 8325-4 frozen stocks were streaked onto brain heart infusion agar (BHI; Millipore 53286, BD Difco 241830) and incubated at 37°C for 24 h to allow colony formation. Overnight cultures were made by inoculating 3 mL BHI broth with three picked colonies and incubating at 37°C at 200 rpm for 18-24 h. Overnight cultures of *S. aureus* strains were sub-cultured in 10 mL BHI broth, grown to log phase (between OD600 = 0.3 and 0.4), and standardized in PBS to 5x10^5^ bacteria/mL for *in vivo* infections. Bacterial concentrations were confirmed by CFU plating.

Mice 6-10 weeks old were anesthetized with ketamine/xylazine solution. A 2 µL suspension of log phase *S. aureus* working stocks were injected into the corneal stroma of mice (approximately 1x10^3^ bacteria/cornea) and mice were euthanized 24 h post infection for examination.

For quantification of corneal opacity, infected corneas were imaged by brightfield then analyzed using ImageJ. Cornea images were converted to greyscale and regions of interest were drawn to exclude areas beyond the cornea. Thresholds were set to exclude areas of the cornea with lighting glare. Mean grey intensity and the area of the cornea were then quantified. The mean grey value of a naïve cornea was subtracted from the values of all infected corneas.

Opacity for a cornea was quantified by multiplying its mean grey value with its area. The maximum opacity was determined by taking the highest mean grey value and multiplying it with the largest area value among the set of infected corneas. Percent opacity of a cornea was determined by dividing its opacity value with the maximum opacity.

For quantification of colony forming units (CFU), whole eyes were excised, homogenized in 1 mL PBS using a TissueLyser II (Qiagen, 30 Hz for 3 min), serially diluted and streaked on BHI agar. Colonies were grown at 37°C for 24 h and counted manually. CFU was also determined immediately after intrastromal injection of *S. aureus* to determine the initial inoculum.

To quantify IL-1β in infected corneas, corneas were excised, homogenized in 500 µL PBS using a TissueLyser II (Qiagen, 30 Hz for 3 min), centrifuged at 16,000 rcf x g for 10 min at 4°C, and clear supernatants were collected for ELISA.

In addition to intrastromal injection of *S. aureus*, a scratch model was performed with the 8325-4 strain. The corneal epithelium of C57BL/6, Myd88^-/-^, and IL-1R1^-/-^ mice were scarified with three central linear abrasions with a 27 Gauge needle (Abbott Laboratories, Chicago, IL), abrasions extended to paracentral cornea. A 5 µL aliquot containing approximately 1x10^6^ CFU 8325-4 strain was applied to the scarified cornea. A sterile trephine (Miltex, Tuttlingen, Germany), was used to make a 2 mm silicone hydrogel contact lens (Night and DayTM, CIBA Vision, Duluth, GA) which was placed over the central abraded corneal to maintain inoculate placement. The contact lenses were removed at 2 h. Mice were euthanized 24 h post infection for examination.

### Histology and immunohistochemistry of infected corneas

Whole eyes were excised, resuspended in 1 mL of David’s Fixative Agent (Poly Scientific R&D Corp, s2250-1GL), and 8-um sections were examined by H&E- and crystal violet-staining, or immunofluorescence staining. A safranin counterstain was not used with crystal violet as it overstained tissues and prevented visualization of crystal violet stained bacteria. For immunofluorescence, neutrophils were detected by incubating cornea sections overnight in rat anti-mouse neutrophil antibody NIMP-R14 (Abcam, 1:100 in 1% FCS-PBS) followed by incubation for 1 h with Alexa Fluor 488-conjugated goat anti-rat antibody (1:250 in 1% FCS-PBS). H&E and gram stains were visualized using a BZ-X710 All-in-One Fluorescence Microscope (Keyence). Corneal thickness was quantified by measuring the pixel length of the scale bar in comparison to the pixel length of infected corneas (from corneal endothelium to epithelium).

### Flow Cytometry

To recover cells from infected corneas, corneas were dissected and incubated at 37°C for 45-60 min in digestion buffer composed of RPMI media (Life Technologies) containing 5 mg/mL collagenase (C0130; Sigma-Aldrich), 0.125 mg/mL DNAse I (QIAGEN 79254), 1% penicillin-streptomycin (Life Technologies), 1% non-essential amino acids (Gibco, 11140-050), 2% HEPES (Gibco, 15630-080), 2% Sodium Pyruvate (Gibco, 11360-070), 10% FBS, and 10 mM CaCl2. Corneas were agitated every 15 min to ensure complete digestion. The samples were transferred to ice and the digestion was stopped by adding FBS equal to 20% of the sample volume. Cells were recovered by filtering samples through cell-strainer capped falcon tubes followed by centrifugation at 300 x g for 5 min and resuspension in flow cytometry buffer.

All subsequent steps were performed at 4°C. Cells from infected corneas, bone marrow, or IP lavage were incubated for 10 min with CD16/32 Ab (BioLegend, Cat# 101320) to block Fc receptors, then incubated for 30 min in the dark with the following antibodies: anti-mouse CD45-FITC (BioLegend, Cat# 103108), anti-mouse CD11b-PerCP-Cy5.5 (BioLegend, Cat# 101228), anti-mouse Ly6G-BV421 (BioLegend, Cat# 127627), anti-mouse Ly6C-PE/Cy7 (BioLegend, Cat# 128017), anti-mouse ADAM10-PE (Biotechne, FAB946P), and fixable viability dye eFluor780 (BD Biosciences). Cells were fixed by incubating for 15 min in the dark with 4.25% paraformaldehyde (BDCytofix/Cytoperm). After cell fixation, cells were resuspended in permeabilization/wash buffer and incubated for 2 h in the dark with anti-mouse IL-1β-APC (Invitrogen, Cat# 17-7114-80) for intracellular cytokine staining. Cells were washed with flow cytometry buffer and quantified using an ACEA Novocyte flow cytometer and NovoExpress software. Neutrophils were identified as CD45^+^CD11b^+^Ly6G^+^ while monocytes were identified as CD45^+^CD11b^+^Ly6G^-^Ly6^+^. The gating strategy is shown in **Fig S2.**

### Computation analysis of α-hemolysin monomers

Structural predictions of α-hemolysin monomers were performed using AlphaFold3 (DeepMind, Google Cloud-hosted implementation). Two constructs were analyzed: the wild-type monomer (UniProt ID: P09616) and a double mutant containing D208E/I275T substitutions. For each construct, AlphaFold3 generated five structural models. The model with the highest overall confidence score (pLDDT and PAE metrics) was selected for subsequent analysis. To assess structural differences, the mutant and wild-type monomer models were compared to each other and to the published structure of the α-hemolysin heptameric pore (PDB ID: 3ANZ).

Structural alignment and root mean square deviation (RMSD) calculations were carried out using the MatchMaker command in UCSF ChimeraX. Structural deviations and conformational differences were examined both quantitatively (via RMSD) and qualitatively through visual inspection.

### Computational modeling of α-hemolysin with ADAM10

To evaluate potential interactions between α-hemolysin and the host cell membrane protein ADAM10, we performed protein-protein complex predictions using AlphaFold3. Input sequences included either the wild-type α-hemolysin monomer (UniProt ID: P09616) or the D208E/I275T mutant monomer, and a single monomer of ADAM10 (UniProt ID: O14672).

Each simulation produced five models, and the model with the highest overall confidence score (pLDDT and PAE metrics) was selected for subsequent analysis. Structural comparisons between the wild-type α-hemolysin-ADAM10 and mutant α-hemolysin-ADAM10 complexes were performed in UCSF ChimeraX. The α-hemolysin components of each complex were superimposed using the MatchMaker tool to assess conformational shifts in the α-hemolysin induced by the mutations. Potential contact sites between α-hemolysin and ADAM10 were also identified by mapping residues within 5Å of the α-hemolysin surface. These residues were color-coded to highlight differences in predicted interaction networks between the wild-type and mutant complexes.

### Statistical Analyses

Experiments were repeated at least twice or with at least 3 biological replicates.

Statistical significance was determined using one-way or two-way ANOVA with either HSD Tukey’s post hoc analysis for *in vitro* studies or by 1-way ANOVA followed by Dunnett’s multiple comparison test for *in vivo* studies. In experiments with only two variables, we used paired or unpaired t-tests. All analyses were performed using GraphPad Prism. Differences were considered significant when the P value was <0.05.

## Acknowledgements

### Author contributions

KL, JTC, JDB, SA, YS and VDJ performed the experiments. CA and MG provided all the clinical strains for this study and AL and RM performed structural analyses on Hla. KL, SA, GRD, MG, RA, CA and EP designed and interpreted the experiments, and wrote the paper.

### Author declarations

The authors declare no conflicts of interest.

### Funding

These studies were supported by NIH grants R01 EY14362 (EP, GRD, RM). The authors also acknowledge departmental support from an unrestricted grant to the Department of Ophthalmology and Visual Sciences from the Research to Prevent Blindness Foundation, New York, NY.

---

Figure S1. USA300 keratitis phenotype is reproducible with 8325-4 keratitis and with topical infection. A-D. A 2 µL inoculum of 5x103 CFU laboratory S. aureus strain 8325-4 was injected into the corneal stroma C57BL/6 or IL-1a/b-/- mice and examined after 24 h. A. Representative images of infected corneas. B. Quantification of viable bacteria in infected corneas by CFU. C. Flow cytometry quantification of neutrophils and monocytes in infected corneas. D. Whole eyes were enucleated, sectioned, and stained histological examination. Representative images are shown of H&E- and crystal violet-staining as well as immunohistochemistry using anti-Ly6G to show infiltrating neutrophils. E. A 5 µL inoculum of 1x106 CFU 8325-4 strain was applied to the surface of abraded corneas from C57BL/6, Myd88-/-, or IL1R1-/- mice and examined after 24 h by immunohistochemistry. Representative images are shown of H&E staining.

Figure S2. Flow Cytometry gating strategy. Representative flow cytometry gating strategy for identification of IL-1β+ neutrophils and monocytes in infected corneas. Total cells from infected corneas were gated on single, live cells and gated for total CD45+ myeloid cells, Ly6G+/CD11b+ neutrophils and Ly6C+/CD11b+ total monocytes.

Figure S3. Rabbit RBC lysis, Inflammasome inhibition and neutrophil elastase and myeloperoxidase (MPO) production A. Conditioned media from USA300, ΔHla, and MRSA clinical isolates were serially diluted prior to incubation with RBCs for 1 h. Intact RBCs were centrifuged and release hemoglobin was assessed by reading optical density. Results are normalized to RBCs treated with triton-x. B-D. LPS-primed BMNs from C57BL/6 were incubated with stimulated for 1 h with 3 mM ATP or 1:10 diluted conditioned media from USA300, ΔHla, and MRSA clinical isolates in the presence of 2 µM NLRP3 inhibitor MCC950, 5 µM caspase-1 inhibitor YVAD, or 10 µM GSDMD inhibitor Disulfiram. Neutrophil Elastase, MPO, and IL-1β in culture supernatatants were quantified by ELISA.

Figure S4. Hla amino acid sequence of USA300 and a CC5 MRSA isolate. Protein BLAST alignment of USA300 and a representative CC5 MRSA isolate showing D208E and I275E amino acid substitutions in the CC5 isolate.

Figure S5. Confidence score predictions for AlphaFold3 models. CC8 (left) and CC5 D208E/ I275T (right) models of monomeric α-hemolysin (top) ADAM10-bound Hla (middle) and Hla-bound ADAM10 (bottom), colored by predicted local distance difference test (pLDDT) per-residue scores. Confidence levels are indicated by color.

Figure S6. Non-specific bands in western blots of MRSA lysates and conditioned media in the absence of primary antibody. Western blot of S. aureus or Pseudomonas aeruginosa whole cell lysates (A) or conditioned media (B) without primary antibodies. Bands in S. aureus wells represent non-specific antibody binding, possibly associated with S. aureus Protein A or Protein G that bind Fc receptors.

Movie S1: D208E/ I275T α-hemolysin resembles active pore structure. AlphaFold3 model of a CC5 α-hemolysin (purple) morphed onto the published structure of an α-hemolysin pore (grey, PDB 3ANZ)

Movie S2: ADAM10 binds D208E/ I275T α-hemolysin differently than CC8 α-hemolysin. The movie shows the ∼90-degree rotation of ADAM10 (grey) when bound to α-hemolysin (green) in a CC8 compared to a CC5 variant. Red: ADAM10 regions within 5 angstroms of CC8 α-hemolysin, Yellow: ADAM10 regions within 5 angstroms of CC5 α-hemolysin, orange: ADAM10 regions within 5 angstroms of both models.

Table S1: MRSA clinical isolates used in this study. The parental USA300 and the ΔHla mutant are from the Nebraska collection; all others are from Mass Eye and Ear (Bispo and Gilmore 2020, Andre, Gilmore 2023) PVL: Panton Valentine leucocidin; ACME: Arginine Catabolic Mobile Element.

## Notes

### Competing Interest Statement

The authors have declared no competing interest.

