## Supplemental figures and tables for "α-hemolysin polymorphisms in methicillin-resistant *Staphylococcus aureus* clinical isolates regulate ADAM10-dependent neutrophil IL-1β secretion"

### Figure S1

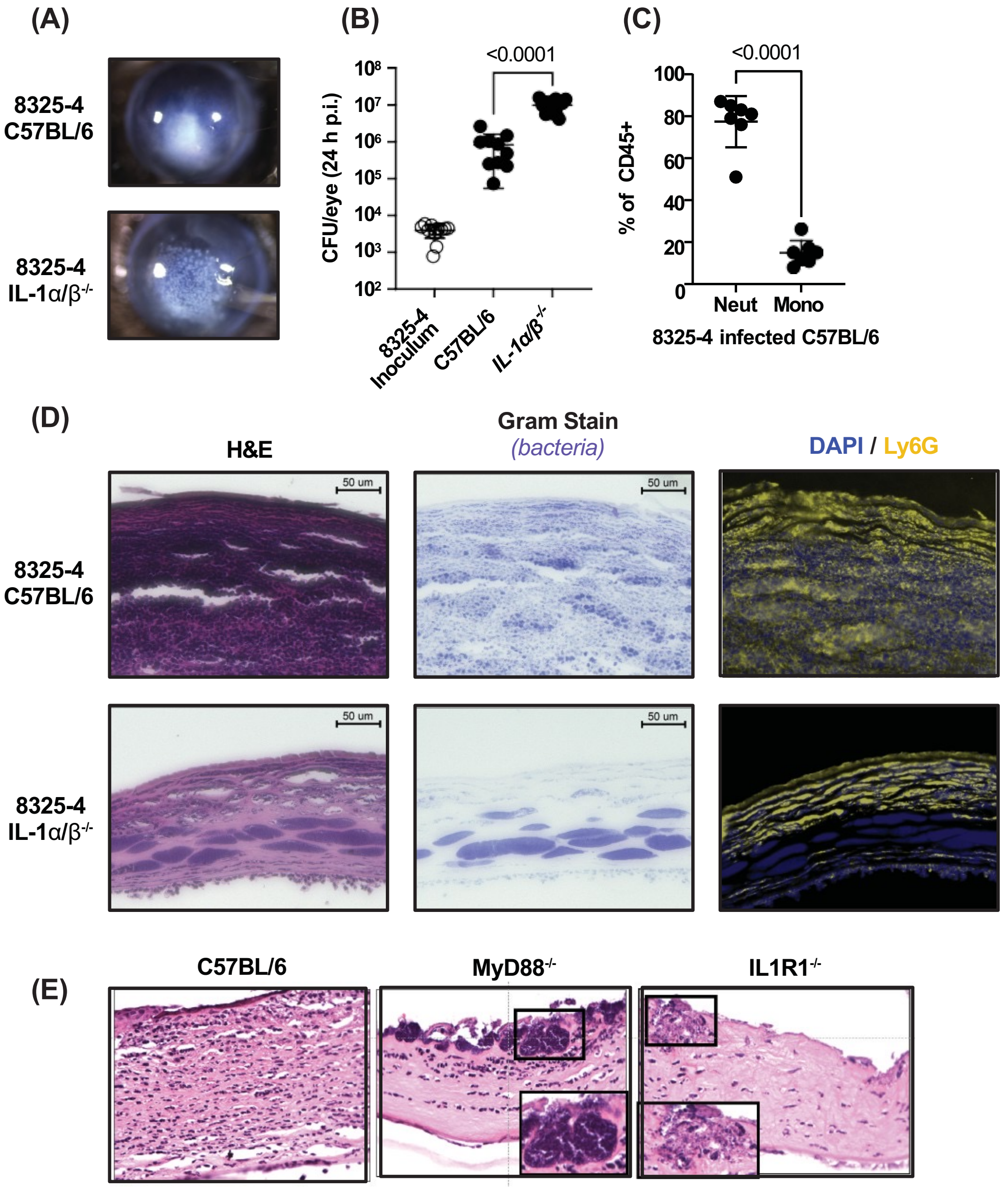

(A)
8325-4 C57BL/6
8325-4 IL-1α/β-/-
(B)
<0.0001
CFU/eye (24 h p.i.)
8325-4 Inoculum
C57BL/6
IL-1α/β-/-
(C)
<0.0001
% of CD45+
Neut
Mono
8325-4 infected C57BL/6
(D)
H&E
Gram Stain
(bacteria)
DAPI / Ly6G
50 um
8325-4 C57BL/6
8325-4 IL-1α/β-/-
(E)
C57BL/6
MyD88-/-
IL1R1-/-

### Figure S2

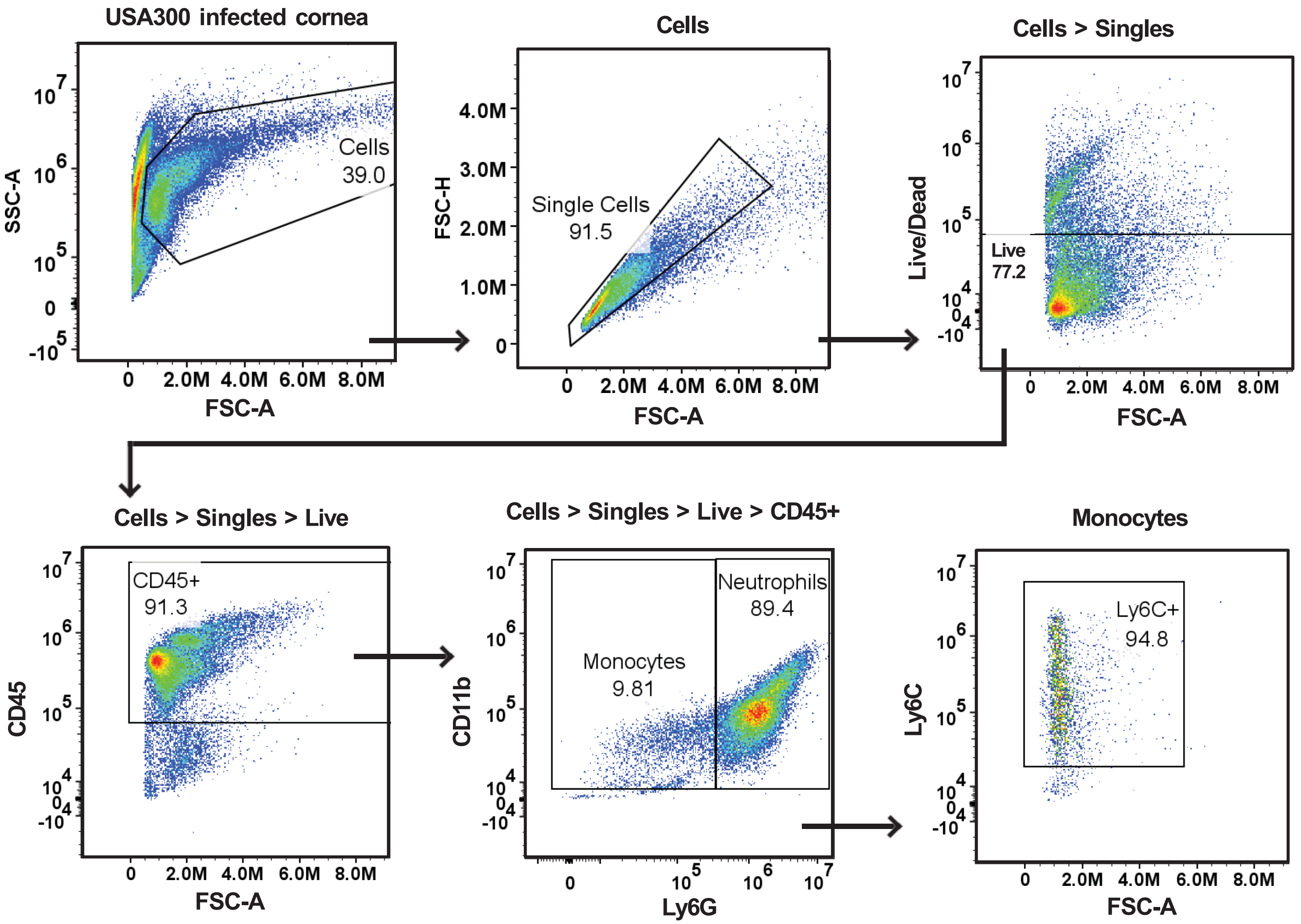
USA300 infected cornea
Cells
39.0
SSC-A
FSC-A
Cells
Single Cells
91.5
FSC-H
FSC-A
Cells > Singles
Live
77.2
Live/Dead
FSC-A
Cells > Singles > Live
CD45+
91.3
CD45
FSC-A
Cells > Singles > Live > CD45+
Monocytes
9.81
Neutrophils
89.4
CD11b
Ly6G
Monocytes
Ly6C+
94.8
Ly6C
FSC-A

### Figure S3

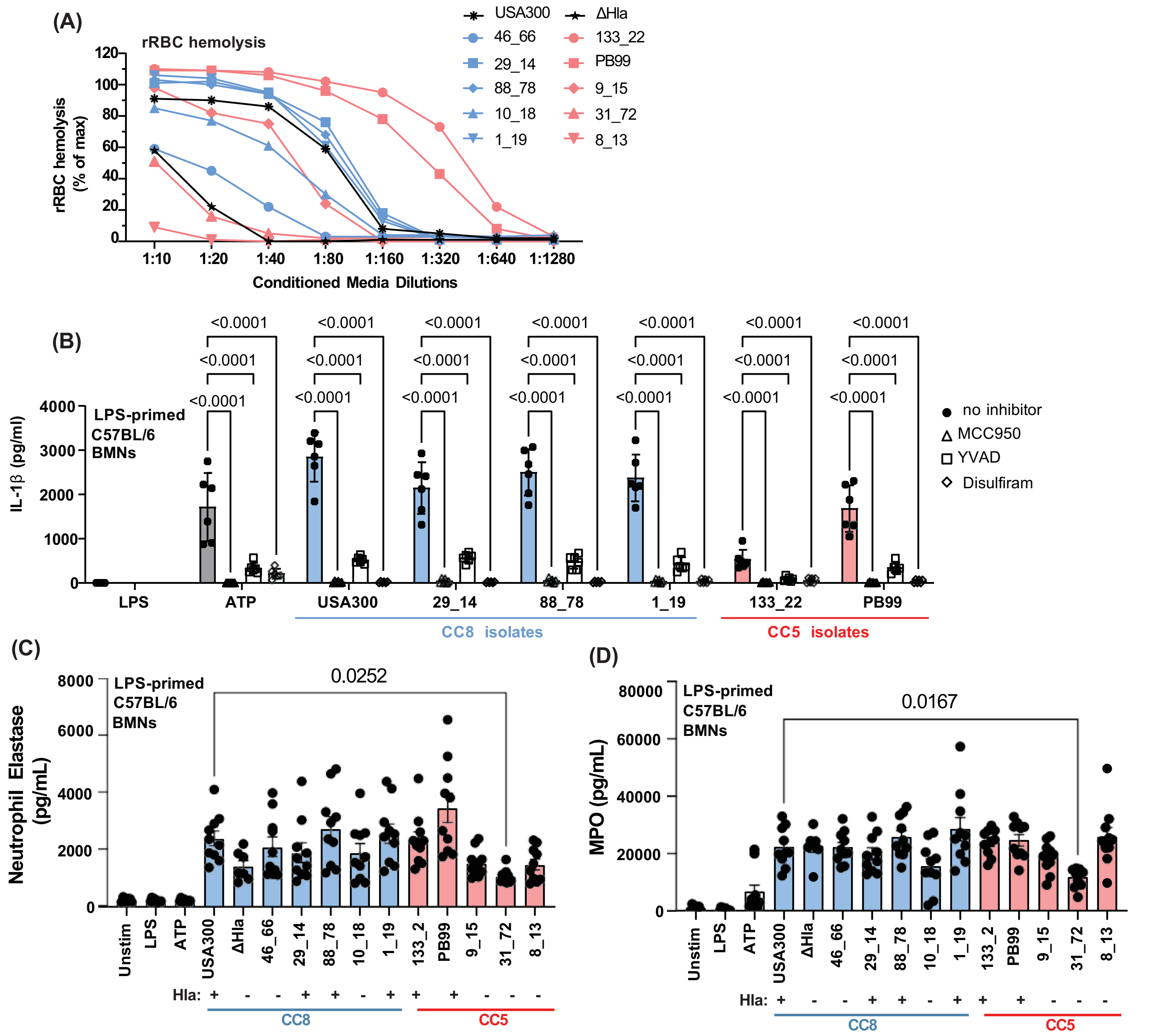
(A)
rRBC hemolysis
rRBC hemolysis (% of max)
120
100
80
60
40
20
0
1:10
1:20
1:40
1:80
1:160
1:320
1:640
1:1280
Conditioned Media Dilutions
USA300
ΔHla
46_66
133_22
29_14
PB99
88_78
9_15
10_18
31_72
1_19
8_13
(B)
LPS-primed C57BL/6 BMNs
IL-1β (pg/ml)
4000
3000
2000
1000
0
<0.0001
LPS
ATP
USA300
29_14
88_78
1_19
133_22
PB99
CC8 isolates
CC5 isolates
no inhibitor
MCC950
YVAD
Disulfiram
(C)
LPS-primed C57BL/6 BMNs
Neutrophil Elastase (pg/mL)
8000
6000
4000
2000
0
0.0252
Unstim
LPS
ATP
USA300
ΔHla
46_66
29_14
88_78
10_18
1_19
133_2
PB99
9_15
31_72
8_13
Hla: + - - + + - + + + - - -
CC8
CC5
(D)
LPS-primed C57BL/6 BMNs
MPO (pg/mL)
80000
60000
40000
20000
0
0.0167
Unstim
LPS
ATP
USA300
ΔHla
46_66
29_14
88_78
10_18
1_19
133_2
PB99
9_15
31_72
8_13
Hla: + - - + + - + + + - - -
CC8
CC5

#### Figure S4

**Alpha-hemolysin Amino Acid Sequence Alignment**

**USA300 (AKR50780.1) - upper sequence Alpha-hemolysin CC5 - lower sequence**

```
ADSDINIKTGTTDIGSNTTVKTGDLVTYDKENGMHKKVFYSFIDDKNHNKKLLVIRTKGT    60
ADSDINIKTGTTDIGSNTTVKTGDLVTYDKENGMHKKVFYSFIDDKNHNKKLLVIRTKGT    60

IAGQYRVYSEEGANKSGLAWPSAFKVQLQLPDNEVAQISDYYPRNSIDTKEYMSTLTYGF    120
IAGQYRVYSEEGANKSGLAWPSAFKVQLQLPDNEVAQISDYYPRNSIDTKEYMSTLTYGF    120

NGNVTGDDTGKIGGLIGANVSIGHTLKYVQPDFKTILESPTDKKVGWKVIFNNMVNQNWG    180
NGNVTGDDTGKIGGLIGANVSIGHTLKYVQPDFKTILESPTDKKVGWKVIFNNMVNQNWG    180

                                   208
PYDRDSWNPVYGNQLFMKTRNGSMKAADNFLDPNKASSLLSSGFSPDFATVITMDRKASK    240
PYDRDSWNPVYGNQLFMKTRNGSMKAAENFLDPNKASSLLSSGFSPDFATVITMDRKASK    240

                                        275
QQTNIDVIYERVRDDYQLHWTSTNWKGTNTKDKWIDRSSERYKIDWEKEEMTN           293
QQTNIDVIYERVRDDYQLHWTSTNWKGTNTKDKWTDRSSERYKIDWEKEEMTN           293
```

Figure S5

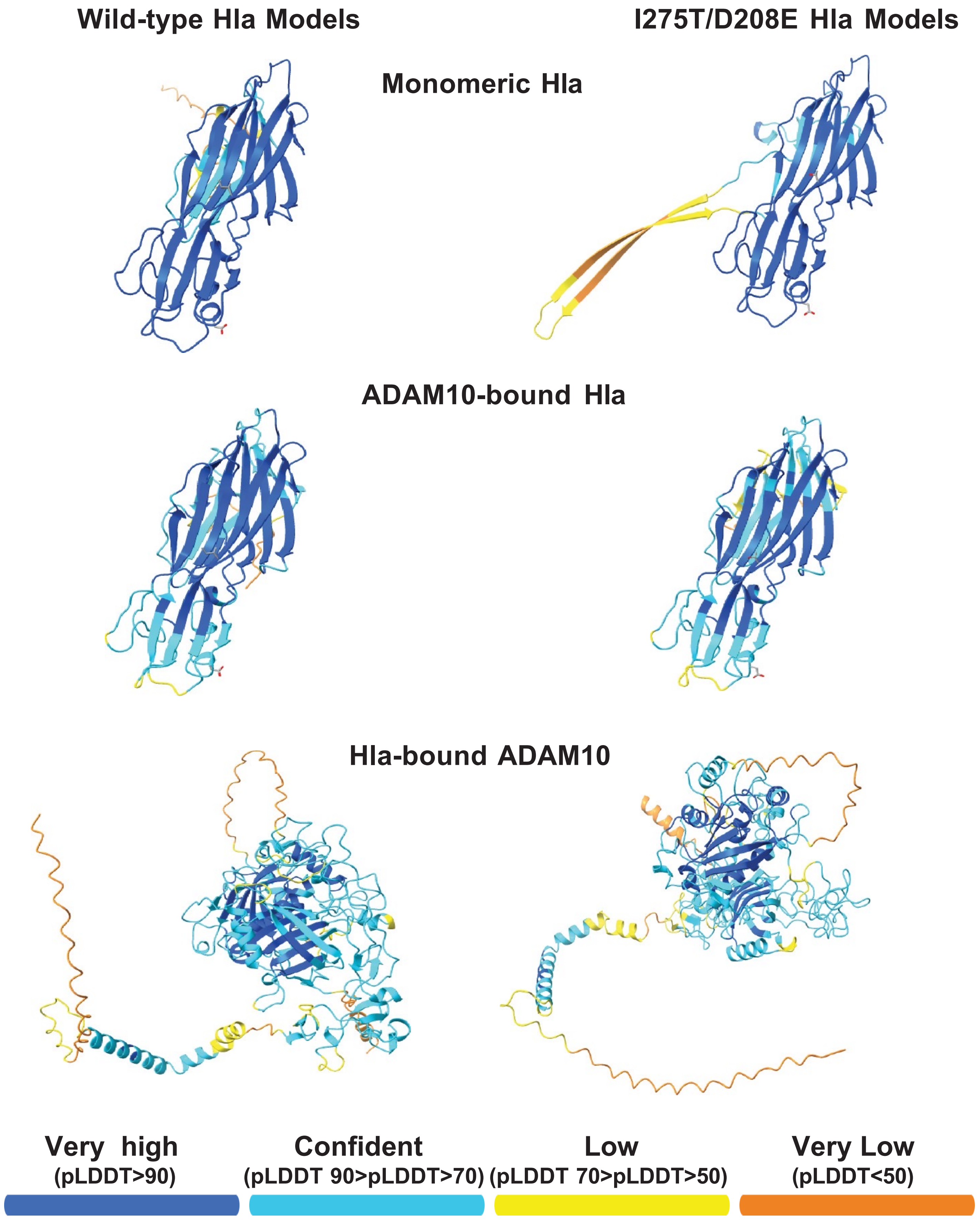

Wild-type Hla Models
I275T/D208E Hla Models
Monomeric Hla
ADAM10-bound Hla
Hla-bound ADAM10
Very high
(pLDDT>90)
Confident
(pLDDT 90>pLDDT>70)
Low
(pLDDT 70>pLDDT>50)
Very Low
(pLDDT<50)

### Figure S6

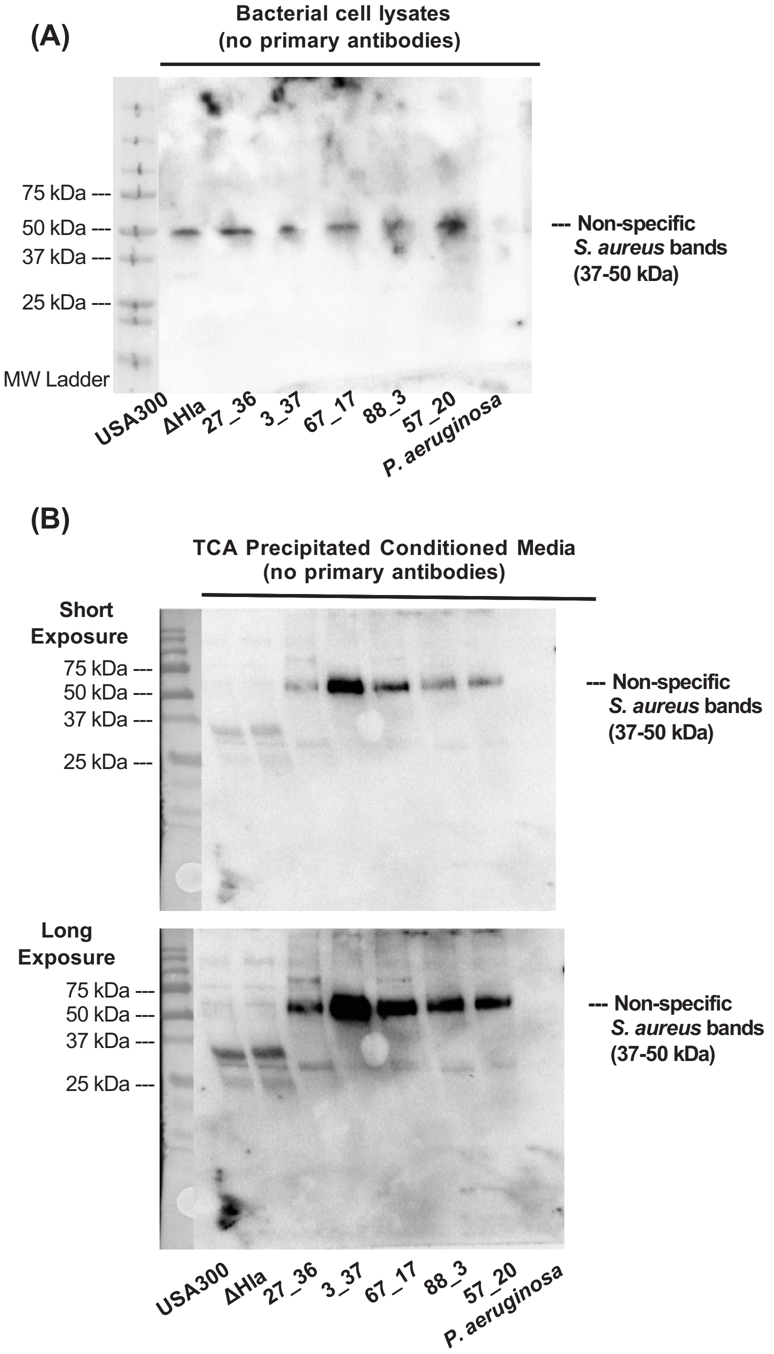

(A)
Bacterial cell lysates
(no primary antibodies)
75 kDa ---
50 kDa ---
37 kDa ---
25 kDa ---
MW Ladder
--- Non-specific
S. aureus bands
(37-50 kDa)
USA300
ΔHla
27_36
3_37
67_17
88_3
57_20
P. aeruginosa
(B)
TCA Precipitated Conditioned Media
(no primary antibodies)
Short
Exposure
75 kDa ---
50 kDa ---
37 kDa ---
25 kDa ---
--- Non-specific
S. aureus bands
(37-50 kDa)
Long
Exposure
75 kDa ---
50 kDa ---
37 kDa ---
25 kDa ---
--- Non-specific
S. aureus bands
(37-50 kDa)
USA300
ΔHla
27_36
3_37
67_17
88_3
57_20
P. aeruginosa

### Table S1

Table S1: MRSA strains used in this study.

| Origins | Isolate ID | CC | ST | SCCmec cassette type | Characterisitics | Infection Demographic | |
|---|---|---|---|---|---|---|---|
| | | | | | | Sex | Age |
| Massachusetts Eye and Ear (doi: 10.3389/fpubh.2020.00204 ) | 46_66 | CC8 | ST8 | IV | PVL+ ACME + | F | 32 |
| | 29_14 | CC8 | ST8 | IV | PVL+ ACME + | F | 26 |
| | 88_78 | CC8 | ST3167 | IV | PVL+ ACME + | M | 28 |
| | 10_18 | CC8 | ST8 | IV | PVL+ ACME + | F | 47 |
| | 1_19 | CC8 | ST8 | IV | PVL+ ACME + | F | 39 |
| | 133_22 | CC5 | ST840 | IV | | F | 86 |
| | PB99 | CC5 | ST1176 | IV | | M | 61 |
| | 9_15 | CC5 | ST5 | IV | | M | 54 |
| | 31_72 | CC5 | ST5 | II | | F | 87 |
| | 8_13 | CC5 | ST105 | II | | F | 88 |
| Nebraska collection (doi: 10.1128/mbio.00537-12) | MG2734 | CC8 | | | MRSA USA300 LAC parental strain | | |
| | NE1354 | CC8 | | | MRSA USA300 a-hemolysin mutant | | |
